# Colour Ambiguity in Real Scenes and the Role of Perceptual Organisation

**DOI:** 10.1101/2021.01.03.423255

**Authors:** Annette Werner, Alisa Schmidt, Julian Hilmers, Lukas Boborzi, Manuela Weigold

**Affiliations:** Max-Planck Institute for Biological Cybernetics, Tübingen, Department for Physiology of Cognitive Processes; University of Tübingen, Institute for Ophthalmic Research

**Keywords:** #thedress, colour ambiguity, colour constancy, segmentation, perceptual organisation, light field

## Abstract

In this study we show a reproduction of the dress-ambiguity phenomenon in a real scene and we report quantitative measurements of the corresponding colour perceptions. The original, real dress, known from #thedress-illusion, was illuminated by combined short- and longwavelength broadband lights from two slide projectors. Test subjects viewing the dress reported to perceive the dress’ fabric and lace colours as blue & black, white & gold or light blue & brown; their corresponding perceptual matches were distributed along the blue/yellow cardinal axis, and exhibited a variability comparable to the ambiguity of the dress photograph. It is particularly noteworthy that the colour ambiguity emerged despite explicit knowledge of the observers about the direction of the light source. Manipulating the background of the real dress (change in chromaticity and luminance, or masking) revealed significant differences between the perceptual groups regarding lightness and colour of the dress. Our findings suggest that observer specific differences in the perceptual organisation of the visual scene are responsible for the colour ambiguity observed for the real dress; in particular, we conclude that colour computations of white & gold viewers focused onto the local region of the dress, whereas the colour processes of blue & black and light-blue & brown viewers were strongly influenced by contextual computations including the background. Our segmentation hypothesis extends existing explanations for the dress’ ambiguity and proposes image based (in the case of the real scene) and high level (in the case of the photograph) neural processes which control the spatial reach of contextual colour computations. The relation between the ambiguity in our real scene and the dress photograph is discussed.

## INTRODUCTION

In 2015, the photography of a dress in a shop (see figure 1) became world famous for uncovering an hitero unseen colour ambiguity: the colours of the dress’ fabric and lace appeared extremely different to different viewers, i.e. either blue and black or white and gold (as it turns out, there are also intermediate percepts, so that it is more precise to talk about a continuum of perceptions). This was independent of the device on which it was viewed, printed or digital.

**Figure 1.**
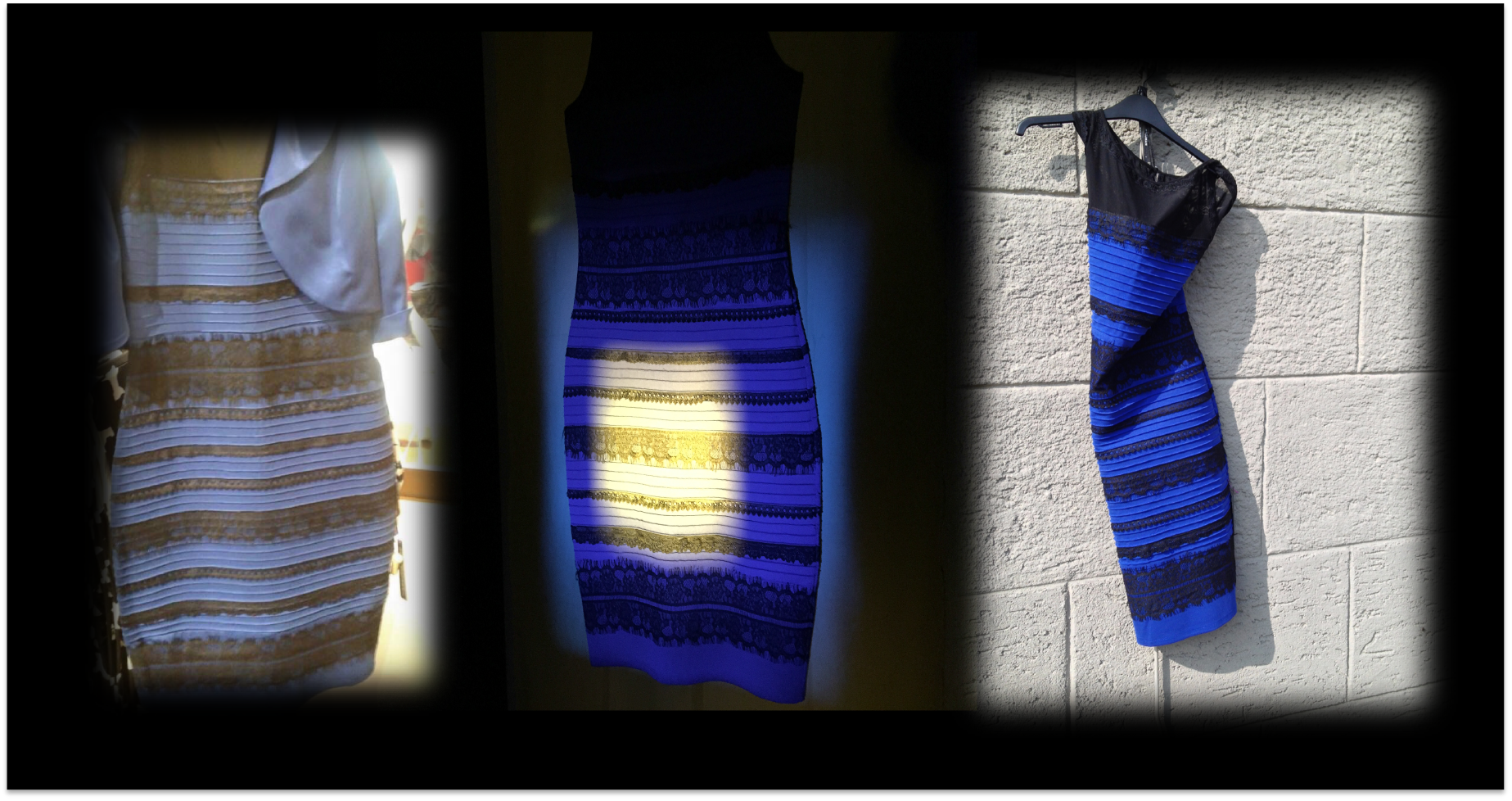
The white/gold or blue/black dress: The widely published original photograph of #the dress (left; source: Tumblr: http://swiked.tumblr.com/post/112073818575/guys-please-help-me-is-this-dress-white-and, photo credit Cecilia Bleasdale) and photographs of the same product [1], taken under different light conditions by one of the authors (A. Werner); middle: a bi-colour presentation of the dress; right: the real dress in real sunlight.

What followed were numerous presentations at public and scientific events and an intensive research into the phenomenon and its underlying causes. For visual neuroscience, the phenomenon is important because like other visual illusions, it offers insights into the neural processes underlying perception. The “dress phenomenon” is however special since it was the first reported case of a colour ambiguity; this is in contrast to other ambiguous percepts in the achromatic domain that are well known since a long time, for example the Necker cube. The colour ambiguity has been a surprise since colour seemed to be always very robust and reliable in the sense that there is in general agreement between subjects (granted normal colour vision) and object colours are recognised even if the illumination and the reflexions from surfaces change (colour constancy). This was not the case with the dress photograph, where viewers strongly disagree. Importantly, the individual differences seem to be specific for the visual scene of the photograph, that is, the dress colour is not ambiguous in general and if seen in another context.

It soon became clear that these differences weren’t simply of semantic origin, that is, that the same colour percept was named differently by different observers. A complicated and yet unexplained relationship seems to exist between the perceptual variation and the age of the observer [2-4], and also – to some extend - with the density of macular pigmentation [5]. Furthermore, several studies have identified the chronotype of the observer as a correlating factor although it remains unclear how these factors may actually cause the ambiguity [3, 4]. Other variations in the sensory or neural equipment of the observers, like pupil size, seem to be an effect rather than the cause for the phenomenon [6]. A corresponding variability of the extension and position of the subjective white point has been proposed but could not fully explain the dress ambiguity [3, 7].

The vast majority of studies, however, has focused on a colour constancy approach, and assume individual differences in the estimation of the illumination in the scene [2-4, 8-14]. The compensation of the illuminant is an integral part of colour constancy, i.e. the reliable encoding of the colour of an object in the face of changing illuminations [15, 16].

In the case of the photograph, it has been argued that ambiguous and sparse information contained in the photograph makes it impossible for the visual sytem to arrive at a stabel and unique solution for the dress colour: namely the uncertainty of the nature of the background scene and the illumination condition, are unclear: it is not clear to a naive observer whether the photograph was taken within a shop, in which case the bright background light would have come from reflexions of the shops illumination in a mirror; alternatively, the photo might be taken outdoors on a shaded balcony with bright sunlight in the background. A distribution of colours along the daylight locus, as in the photograph, could make this task even more difficult [17, 18].

For resolving this problem, it has been suggested that the visual system could use top down information, such as priors drawn from the first (random) perceptual outcome of the neural colour code, that is, the colour perceived when seeing the dress photograph for the first time [19]; alternatively, it could rely on previous experiences in order to make specific assumptions about the nature and properties of the illumination, such as its spectrum or the number and position of light sources [2-4, 7, 9-13, 19, 20]. For example, the popular sun/shadow hypothesis (as demonstrated in figure 2) states that subjects who perceive the dress as blue & black (BB) assume, the dress to be in bright sunlight and their compensation processes therefore intend to “make” the dress darker and more blue; white & gold (WG) viewers on the other hand, are thought to compensate an assumed shadow on the dress and therefore “make” the dress brighter and more whitish. A contribution from brain regions involved in higher cognition has been further substantiated by a study using magnetic resonance imaging (fMRI), which found in the frontal and parietal brain areas of WG viewers a higher activation in response to viewing the dress photograph than in BB viewers [21].

**Figure 2.**
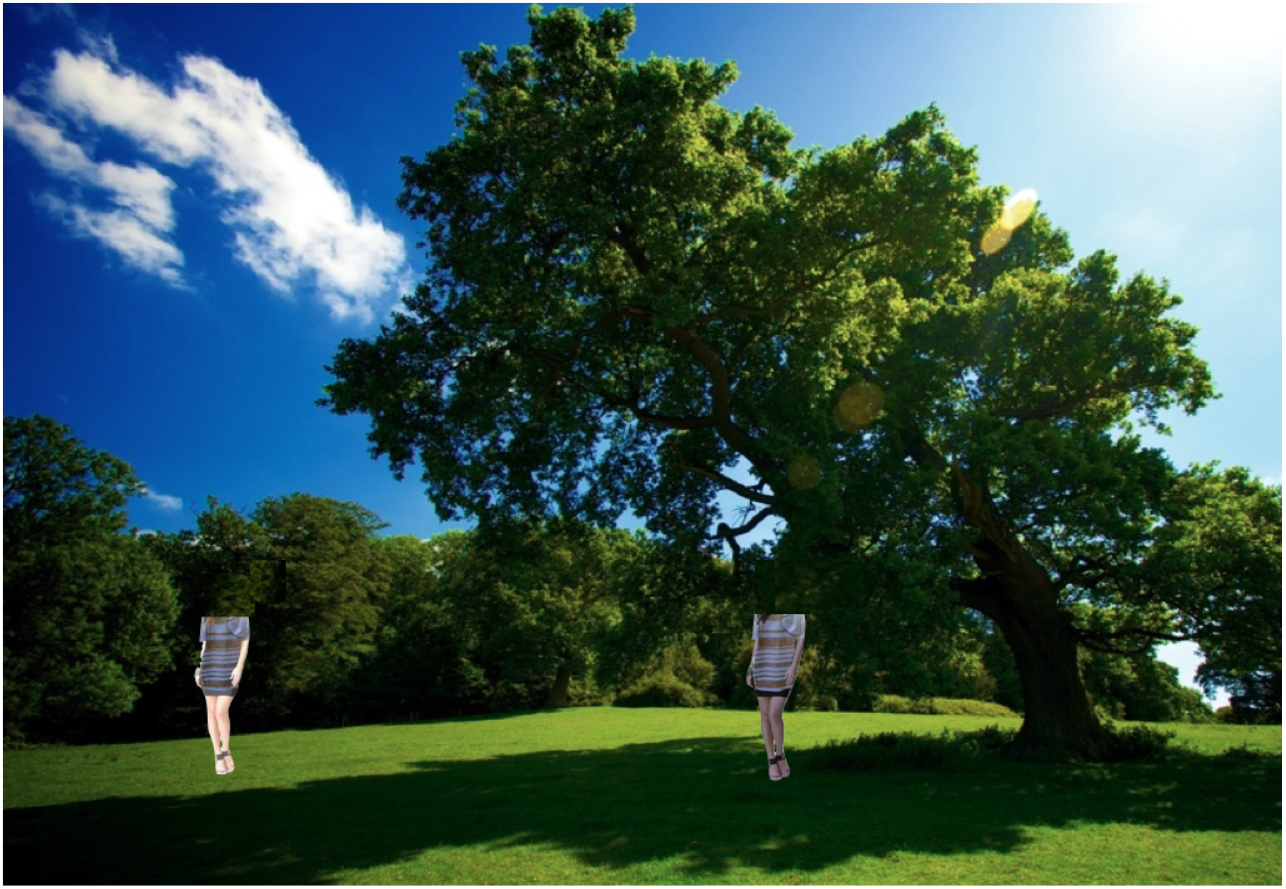
Illustration of the sun/shadow hypothesis, modified after the demo by Yukiyasu Kamitani. Note that the figures’ dress is identical in both conditions, sun and shadow, but the values for the ladys skin are not, i.e. they are darker in the shadow figure, and brigther in the sun figure; in other words, the illustration does not reproduce an effect of different illuminations on both figures but rather the effect of chromatic induction. In any case, it demonstrates nicely the different colour percepts.

Our understanding of colour ambiguities were further challenged by the discovery of another ambiguous photograph of a shoe - “the shoe illusion” – where colours are not distributed along the blue/yellow (i.e. daylight) axis, but rather spread roughly perpendicular to it, along the red/green cardinal axis [22-24]. Furthermore, it has been recently demonstrated that colour ambiguity is not unique for the dress photograph after all, but can also be produced by other, artificial images of objects, provided they exhibit a particular combination of colours and require a certain amount of scene interpretation from the observers [23, 25].

But does colour ambiguity exist in real scenes? So far, all of the above cited studies focused on the original dress photo, or virtual scenes. A reproduction of the ambiguity in a real scene would open the door to study more directly the peculiarities of the underlying colour computations. Indeed, a reproduction of the dress ambiguity with the real dress was presented at two public events: one during the Wellcome Collection’s “On Light” event 2015 [26] and another during the Vision Science Society (VSS) Demo Night 2015 [27, 28]. In these two independent demonstrations, the real dress (blue version, RomanOriginals©) was presented under bi-colour illuminations (blue and yellow, LEDs or filtered broadband light from halogen lamps). However, these were mass demonstrations, and the observers responses were not evaluated in detail.

Here we show, to our knowledge for the first time, quantitative measurements of the reproduced ambiguous percepts in a real scene with the original real dress. Importantly, in the present study we also manipulated the chromatic context of the dress: this follows from anecdotical reports during our previous demonstrations which indicated strong contextual influences on the perceived colour of the dress when seeing a human figure next to it [28]. Our findings suggest an important role for the dress’ background in the emergence of the ambiguity and we conclude that specific differences in the perceptual organisation and consequently the processing of contextual information between BB and WG viewers are responsible for the observed colour ambiguity of the dress, even in the absence of an ambiguous light situation.

## METHODS AND MATERIAL

### Subjects

26 subjects participated in the experiments; they were selected according to their a priori statements on their perception of the dress colour in the photograph (figure 1, left side); the classification of the subjects was confirmed in our study by their matches of the dress in the photograph (figure 5 shows the approximate appearance of these matches). In the following, the subjects will be referred to as the white-and-gold group (WG; n=11, average age 24.5 years ± 0.8, 4 males, 7 females), the blue-and-black group (BB; n=11, average age 24.1 years ± 0.8, 5 males, 6 females) and the light-blue-and-brown group (LB; n=4, average age 24.0 years ± 0.4, 2 males, 2 females).

### General visual performance

Prior to the main experiments, colour discrimination (Cambridge Colour Test; Cambridge Research Systems), and visual acuity [29] were tested; all subjects had normal colour vision or normal or corrected-to-normal visual acuity.

### Experimental setup and stimuli

In the experiments, the real dress and a printout of the original photo were presented in a laboratory room which was either illuminated by ceiling light or by the light of two slide projectors. Figure 3 shows the arrangement of photo, dress, and slide projectors during the experiments. Colorimetric data of the stimuli were obtained from the measurements with a calibrated spectrometer (CAS140CT-152, Instrument Systems). Colour loci and spectral characteristics of the real dress, the dress photograph and the background cloths and their illuminants are despicted in figure 4.

**Figure 3.**
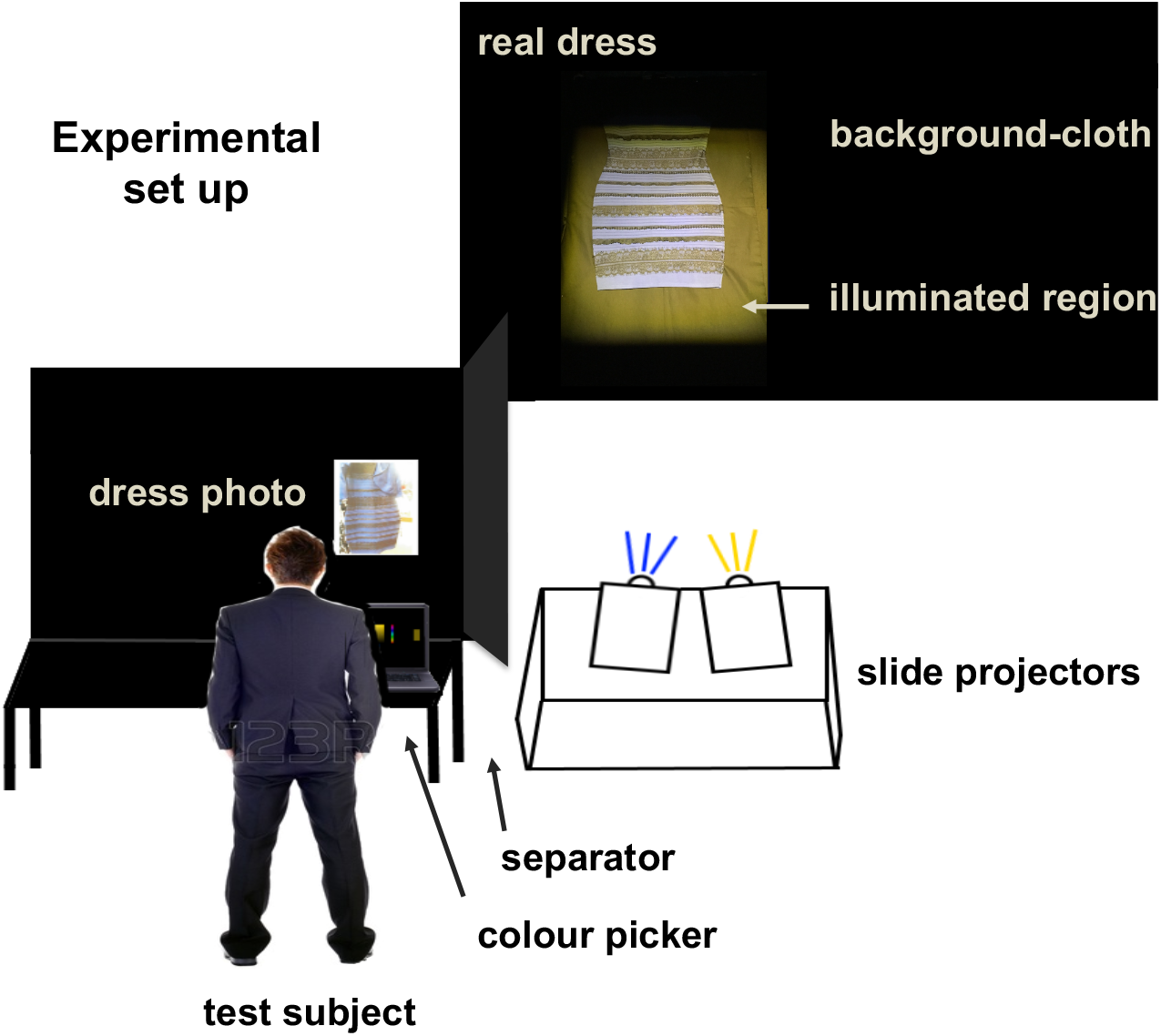
Scheme of the experimental set up. The real dress (upper right) was placed on either a black (shown here) or a yellow background cloth, and was illuminated by the light of two slide projectors. The printout of the photo (lower left) was presented on a vertical black cardboard background, and was shielded to the side by a black cardboard separator. Please note that the stimuli (real dress <u>or</u> dress photo) were present only one at a time during the experiments; subjects made colour matches using a custom made colour picker program.

**Figure 4.**
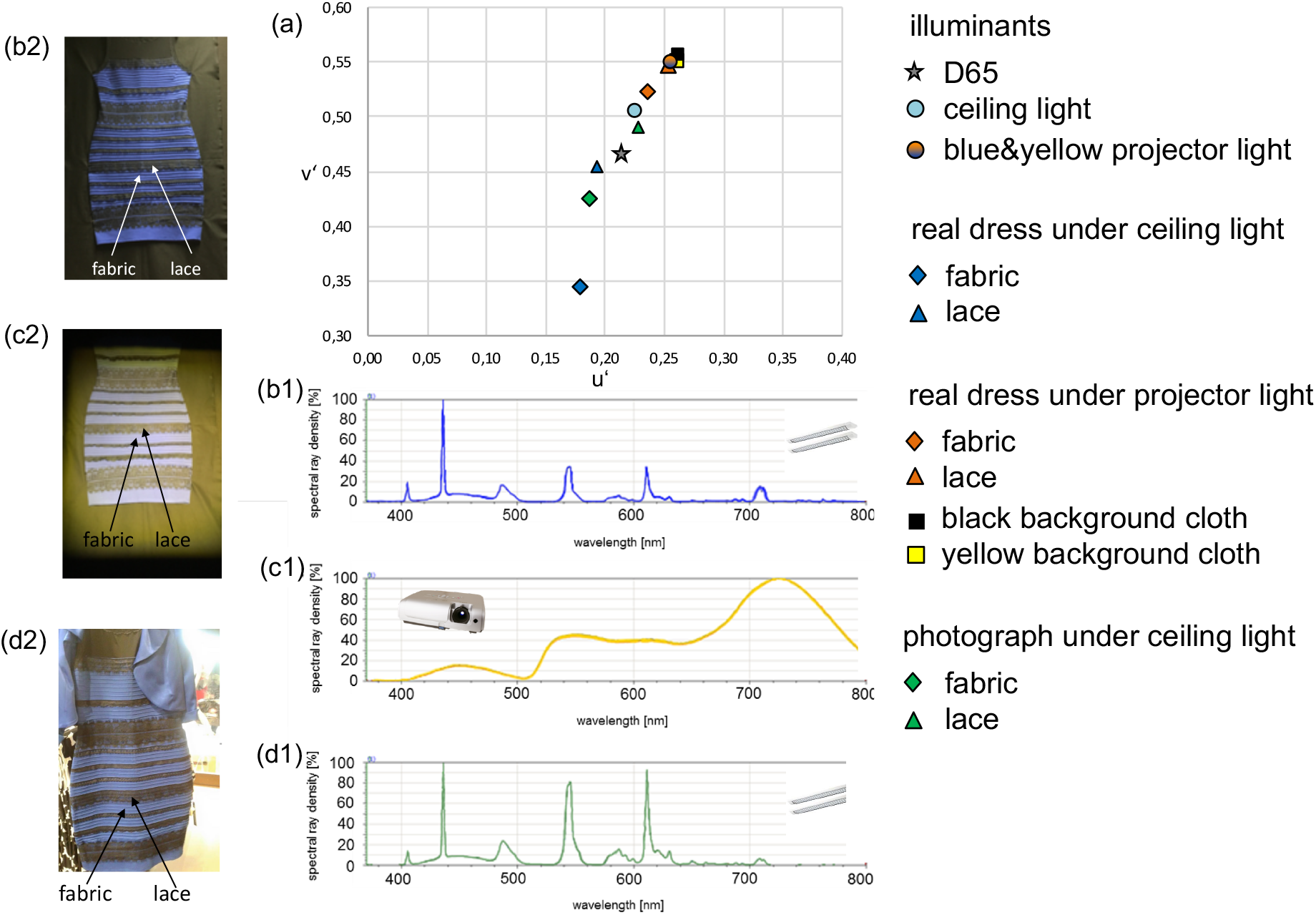
Chromaticities and spectral characteristics of the dress in the photograph and the real dress and their illuminants. Figures on the left show the real dress in different light conditions and the dress photograph. (a) CIE1976 UCS chromaticity diagram with corresponding colour loci, for the symbols see inserted text; (b1) spectrum of the fabric and (b2) image of the real dress under ceiling light; (c1) spectrum of the fabric and (c2) image of the real dress under projector light; (d1) spectrum of the fabric and (d2) image of the printout of the dress photograph, presented under ceiling light; arrows indicate the regions (fabric and lace) from which the measurements and colour matches were obtained.

### Experiments using the dress photograph

The original photo (http://swiked.tumblr.com/post/112073818575, accessed on April 27^th^ 2015), was printed out in A4 format on Plano^®^Speed Business Copy Paper (Papyrus) using a colour laser printer (Brother DCP-9045CDN). The printout of the photo was presented vertically on a black cardboard background, illuminated from above by the labs ceiling light (Osram Lumilux Cool White FQ 49W/840 HO, 4000 K). The observers viewed the photograph at eyelevel from a distance of 60 cm, in two conditions: 1) the full view condition: the photograph was shown as originally posted (size 20 × 30 cm); and 2) a masked condition: only two small regions (each 1 × 1 cm) of the dress were visible, one showing a patch of the fabic, the other showing a patch of the lace), the remainder of the photo being covered by black cardboard. Figure 4 shows the chromaticities of fabric and lace of the dress in the photograph (fabric: u’ = 0.1859, v’ = 0.4262, L = 14.82 cd/m^2^; lace: u’ = 0.2269, v’ = 0.4915, L = 6.26 cd/m^2^).

### Experiments using the real dress

The original real dress (blue version) was purchased from the company RomanOriginals© (Brimingham, UK), its material consists of 68% viscose, 27% polyamide, 5% elastane. The subjects viewed the dress from a distance of 300 cm (figure 3). For simulating the visual scene in the photo, the dress was presented hanging against one of the labs walls, which was covered by either a black or a yellow cloth (149 × 95 cm; cotton-jersey mix; their chromaticities are despicted in figure 4; note that the two backgrounds are very similar in hue and chroma, and differ mainly in luminance (black cloth: u’ = 0.2603, v’ = 0.5507; L = 6.824 cd/m^2^; yellow cloth: u’ = 0.2623, v’ = 0.5570, L = 202.286 cd/m^2^).

There were three presentation- and test-conditions: (1) the real dress illuminated by projector lights: the dress was presented in a dark room, whereby the light from two slide projectors (Leica Pradovit P600) illuminated the skirt part of the dress (w × h = 45 × 50 cm) horizontally from the front (see figure 3; fabric: u’ = 0.2351, v’ = 0.5240, L = 12.0 cd/m^2^; lace: u’ = 0.2521, v’ = 0.5475, L = 8.3 cd/m^2^); the total area illuminated by the projectors was 70 × 95 cm (w × h) and extended into the immediate background. The projectors were placed next to each another on a stand (hight 60 cm) with a distance of 280 cm from the dress. The projectors light sources were halogen lamps (Osram 24V/2450W), in one of the projectors filtered through a yellow glass filter (λ_max_ 530 nm, 2 mm, Schott), in the other filtered through a blue gel filter (λ_max_ 460 nm, Kodak Wratten gelatin filter CAT 1707173 No.34). The lights themselves were not visible to the observers, the chromaticities of the projectors illumination are despicted in figure 4. (2) the masked condition: for the tests involving the masked paradigme, the illuminated region (w × h = 20 × 30 cm) of the dress was restricted to the central part of the skirt with the help of an aperture slide. 3) the real dress illuminated by ceiling light: the dress was presented in a fully lit room, whereby the illumination was provided from above by the labs ceiling light (same as the one illuminating the dress photograph; fabric: u’ = 0.1774, v’ = 0.3463, L = 2.53 cd/m^2^; lace: u’ = 0.1929, v’ = 0.4556, L = 0.369 cd/m^2^).

### Colour matches using a colour picker program

The subjects were instructed to match, using a custom made colour picker program, the appearance of the dress fabric or lace with that of a sample patch (5 × 2 cm), which was presented on a labtop screen (Sony Vaio VPCEC4M1E) in front of them (figure 3). The sample patch was surrounded by black cardboard and subjects wore a black cloak over their clothes in order to avoid reflexions of their clothing to interfere with the matches displayed on the laptops screen.

### General procedure

After testing the general visual performances of the subjects, the actual experiments started with the presentation of the real dress illuminated by projector light, first on a black background, then on a yellow background. The subjects were asked to use the colour picker and to make colour matches of indicated areas of the dress (see arrows in figure 4); after each test the subjects were asked to name the colour they had matched; the matching chromaticities and the corresponding colour names were both recorded. The experiments continued with the presentation of the photo under celling light, where subjects made again colour matches for indicated positions on the photo and then named the matching colours. Only at the end of the experimental session, observes were shown the real dress under ceiling light, and matched the appearance of fabric and lace with the colour picker.

### Data analysis and statistics

The observers colour matches were specified in CIE 1976 UCS chromaticiy coordinates u’v’, and L (cd/m^2^) for the lightness match. For determining within-observer perceptual changes between conditions, we calculated separately the euclidian distance (Δu’v’) between the respective colour loci with the u’v’ coordinates (u’1 v’1) and (u’2 v’2) and the differences (ΔL) of the corresponding luminance values (L1 and L2) of the observers matches by:

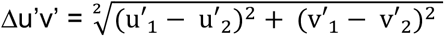

and

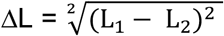

Statistical analysis of the data was carried out using the non-parametric Mann-Whitney-U test (2-tailed) for group differences and the 1-sample Wilcoxon-Rank test (2-tailed) for comparing the obsevers matches with the colourimetric coordinates; the significance level was set at 0.05.

## RESULTS

### 1. #THE DRESS PHOTOGRAPH

This set of experiments aimed at bringing the perceptual settings of our group of observers into context with the findings from earlier studies on the dress photograph. Please note that these measurements were taken <u>after</u> concluding the experiments with the real dress but are shown here first in order to allow a better evaluation of our findings in the real scene experiments. Figure 5 presents the appearances of the colour matches made by BB, WG and LB viewers for the fabric and lace in the photograph.

**Figure 5.**
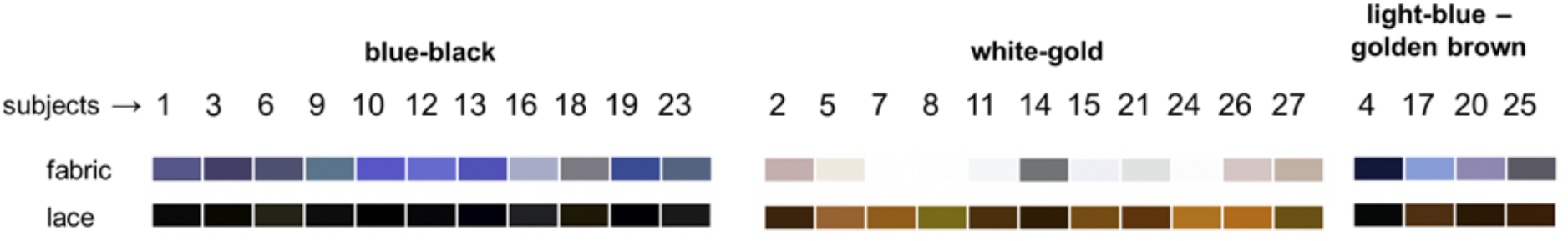
Colours matched by BB, WG and LB viewers for the fabric and lace of the dress in the photograph; please note this reproduction can only be an approximation of the appearance of the actual matching samples.

Figure 6 shows the colour loci of the settings made by the observers for the fabric as well as the lace of the dress in the photo. The individual matches were consistent with the a priori classifications of the observers, but the individual perceptions varied within each group. Altogether, the settings for the fabric were distributed along the blue/yellow cardinal axis in colour space, extending from a region near the colour locus of the fabric (and near D65) into the blue region of the colour diagram; the settings of WG viewers were located between the achromatic region (near D65) and the colour locus of the fabric (i.e. a desaturated blue; WG: u’_mean_ = 0.198, v’_mean_ = 0.461; fabric: u’ = 0.1858; v’ = 0.4262) while BB and LB chromaticity settings for the fabric were located in the blue region (BB: u’_mean_ = 0.183, v’_mean_ = 0.349; LB: u’_mean_ = 0.189, v’_mean_ = 0.372). The distribution of the settings along the blue/yellow axis is mirrored in the variation of the v’ coordinates, which differed significantly between WG viewers and BB viewers (p_WG-BB_ < 0.0001) and LB viewers (p_WG-LB_ = 0.001), respectively; there was no difference between BB and LB viewers (p_BB-LB_ > 0.5).

**Figure 6.**
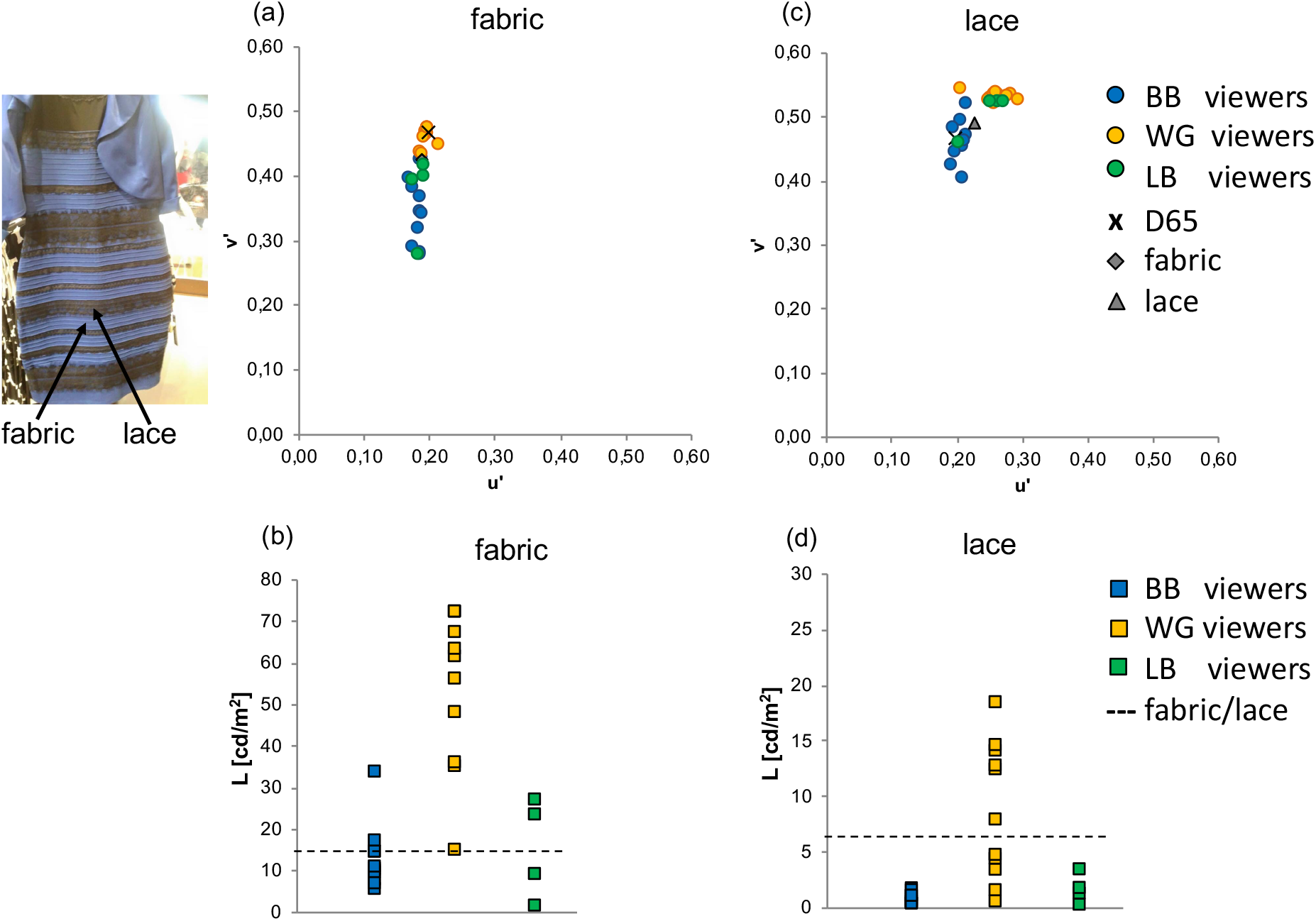
Individual chromaticities and luminance settings for the appearance of the dress in the photograph. Datapoints are coloured according to the a priori self-classification of the subjects as BB, WG or LB viewers; x denotes the colour locus of D65, diamond and triangle mark the colour loci of fabric and lace, respectively; the dotted lines indicate the luminance of the fabric or lace. (a) u’v’ chromaticities and (b) luminance of the matches for the fabric; (c) u’v’ chromaticities and (d) luminance of the matches for the lace, respectively. The image on the upper left shows the original photograph of the dress; arrows indicate the regions (fabric and lace) from which the measurements and colour matches were obtained.

Concerning lightness, the matches of WG viewers were significantly higher (L_mean_ = 54.32 cd/m^2^) than the actual luminance of the dress (L_fabric_ = 14.82 cd/m^2^, p_WG-fabric_ = 0.002), and the matches of BB (p_WG-BB_ < 0.0001) and LB viewers (p_WG-BB_ = 0.006), respectively. The matches of BB viewers (L_(mean)_ = 12.65 cd/m^2^) and LB viewers (L_mean_ = 15.05 cd/m^2^), on the other hand, did not differ significantly from the actual luminance of the fabric and one another (p_BB,LB-fabric_ > 0.1).

For the lace, the chromatic settings showed remarkable individual differences, too: the settings of WG and LB viewers were near its colour locus in the yellow-orange (“gold-brown”) region (WG: u’_mean_ = 0.263, v’_mean_ = 0.529; LB: u’_mean_ = 0.247, v’_mean_ = 0.507); the settings of BB viewers, on the other hand, were in or near the achromatic region (u’_mean_ = 0.205, v’_mean_ = 0.460). v’ coordinates differed significantly between all groups (p_WG-BB_ > 0.0001; p_WG-LB_ = 0.04; p_BB-LB_ = 0.026), in addition, WG and BB viewers also differed in their u’ coordinates (p_WG-BB_ < 0.0001). The lightness matches of BB viewers for the lace were significantly lower than the actual luminance of the lace (BB: L_mean_ = 0.70 cd/m^2^; p_BB-lace_ = 0.002); WG and LB viewers settings were on average close to the photometric luminance (WG: L_mean_ = 8.15 cd/m^2^, p_WG-lace_ > 0.1; LB: L_mean_ = 1.69 cd/m^2^, p_LB-lace_ > 0.1).

#### Summary and conclusion of part 1

Overall, the chromaticity settings of the observers corresponded to their a priori statements as WG, BB or LB viewers. Furthermore, the distribution of their perceptual matches for fabric and lace of the dress is consistent with the findings reported in previous studies of the dress photograph [2, 3, 8]; the alignment of the settings along the daylight locus confirms the notion that there is a continuum of perceptions rather than a sharp clustering in three groups, as may be suggested from the colour naming categories. Nevertheless, the v’ chromaticity coordinates differed significantly between the WG viewers on one hand and the BB and LB viewers on the other hand, indicating a perceptual ambiguity. In addition, an ambiguity was also observed in the lightness domain: relativ to the actual luminance of the dress, WG viewers overestimated the value of the fabric, whereas BB viewers underestimated the luminance of the lace.

### 2. REPRODUCTION OF THE AMBIGUITY PHENOMENON IN REAL VISUAL SCENES

#### 2.1. The real dress on “black” background

The real dress was first presented against the black background cloth, and illuminated by a mixture of blue and yellow light from the two slide projectors. When naming the perceived colours, the majority of the observers (20 of 26) described the colour of the dress as white with a slight colour tint (whitish or very desaturated blue), and the lace as dark-bronze or gold-coloured; however, when asked to make an exact match of the colours of the dress fabric, a more differentiated picture emerged (see figures 7 and 8).

**Figure 7.**
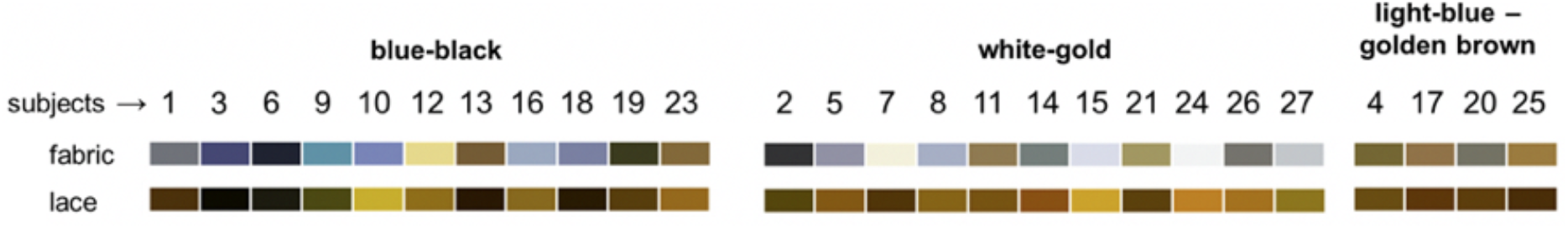
Appearance of the colour matches made by BB, WG and LB viewers for the real dress on black background.

**Figure 8.**
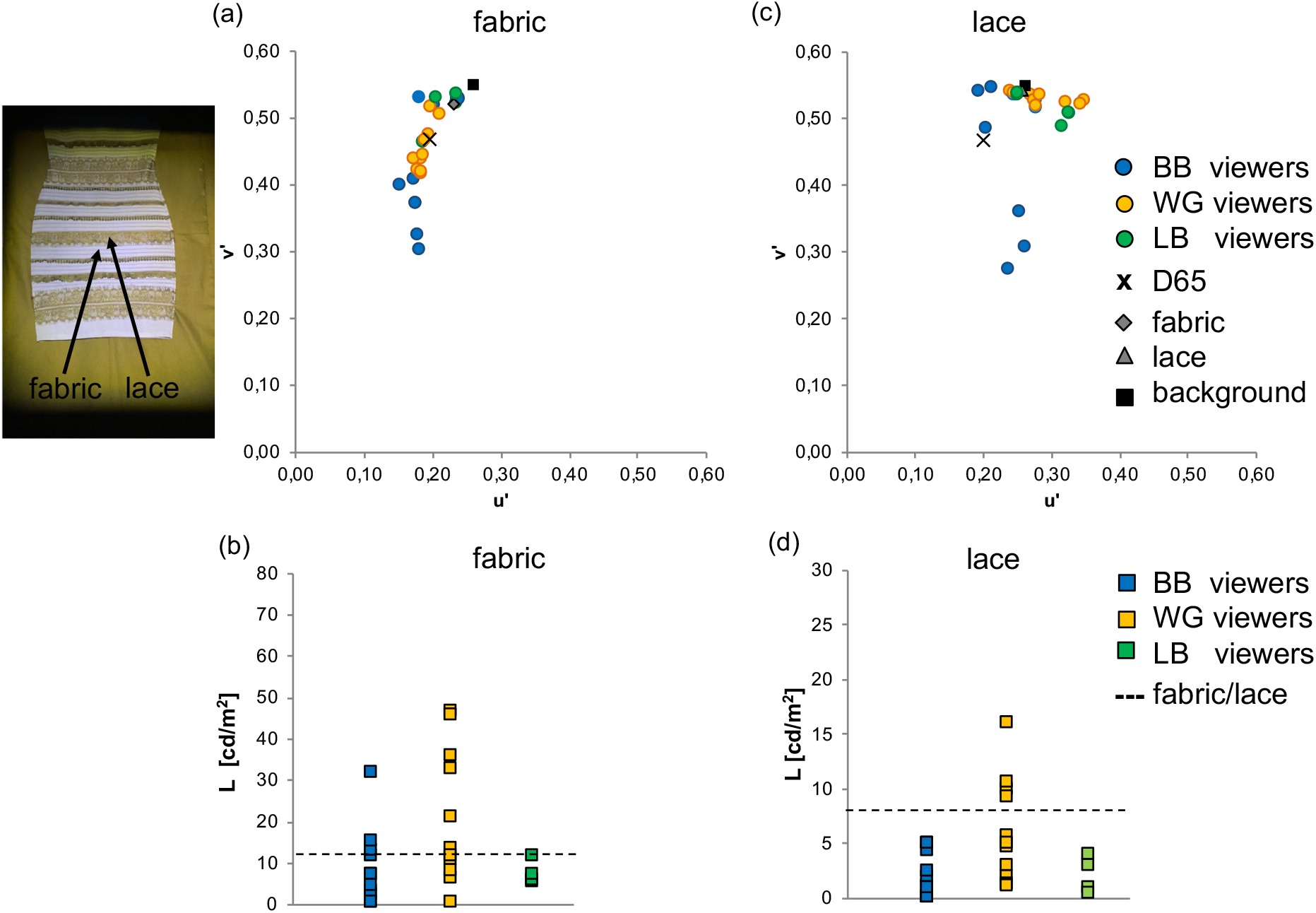
Individual chromaticities and luminance settings for the appearance of the real dress fabric (left) and lace (right), presented on a black background cloth under projector light. (a) u’v’ chromaticities and (b) luminance of the matches for the fabric; (c) u’v’ chromaticities and (d) luminance of the matches for the lace, respectively. Inserted image on the upper left shows the real dress on the black background as presented in this experiment; arrows indicate the regions (fabric and lace) from which the measurements and colour matches were obtained. Same symbols as in figure 6; black square marks the colour locus of the background cloth.

As can be seen in figure 8, a considerable variability was observed for the v’ coordinates, i.e. along the blue/yellow cardinal axis, although the matches did not differ significantly between observers: the settings of all WG viewers were close to D65, i.e. achromatic or highly unsaturated yellowish or bluish (WG: u’_mean_ = 0.190; v’_mean_ = 0.454); the LB viewers were located in the same chromatic region, i.e. within the cluster of WG viewers (LB: u’_mean_ = 0.218; v’_mean_ = 0.511). Notably, the settings of four BB viewers were also within the region of WG viewers, the remaining BB viewers located their settings in the blue colour region (u’_mean_ = 0.191; v’_mean_ = 0.427). Lightness matches of all groups of observers (WG: L_mean_ = 21.03 cd/m^2^; BB: L_mean_ = 8.69 cd/m^2^; LB: L_mean_ = 7.42 cd/m^2^) did not differ significantly from the photometric luminance of the fabric (p_(WG, BB, LB)-fabric_ > 0.1), but settings of WG viewers were significantly higher than those of BB viewers (p_WG-BB_ = 0.047).

For the lace, we observed no significant differences between the perceptual groups, neither in their chromaticity nor in their lightness settings. With the exception of three BB viewers, whose matches were located in the blue region, all other matches were located near the lace’ colourimetric locus in the yellow-orange region (lace: u’_mean_ = 0.262, v’_mean_ = 0.534; L_mean_ = 2.94 cd/m^2^), and were consistent with the colour names „gold“ or „brown (WG: u’_mean_ = 0.262, v’_mean_ = 0.534, L_mean_ = 6.19 cd/m^2^; BB: u’_mean_ = 0.220, v’_mean_ = 0.420, L_mean_ = 3.87 cd/m^2^; LB: u’_mean_ = 0.254, v’_mean_ = 0.428, L mean = 2.02 cd/m^2^).

#### 2.2. The real dress on yellow background

After the experiments using a black cloth as background, the dress was presented on a bright yellow cloth. Following the background change, observers reported a perceptual shift towards blue, which was accompanied by a reduction in the perceived luminance; this was also mirrored in the colour matches (figures 9 and 10).

**Figure 9.**
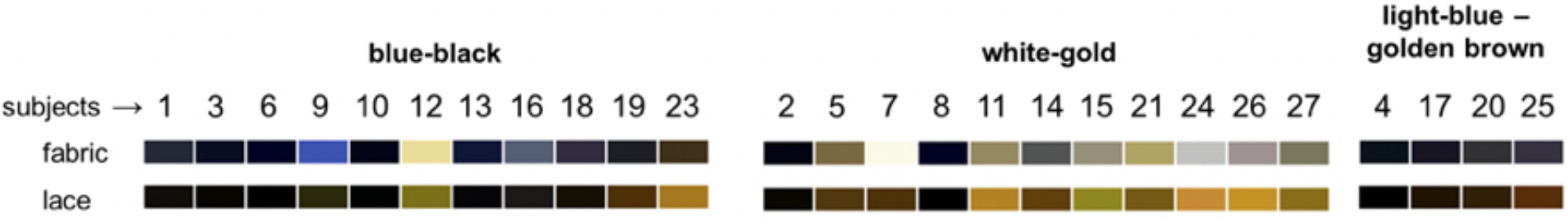
Appearance of the colour matches made by BB, WG and LB viewers for the real dress on yellow background.

**Figure 10.**
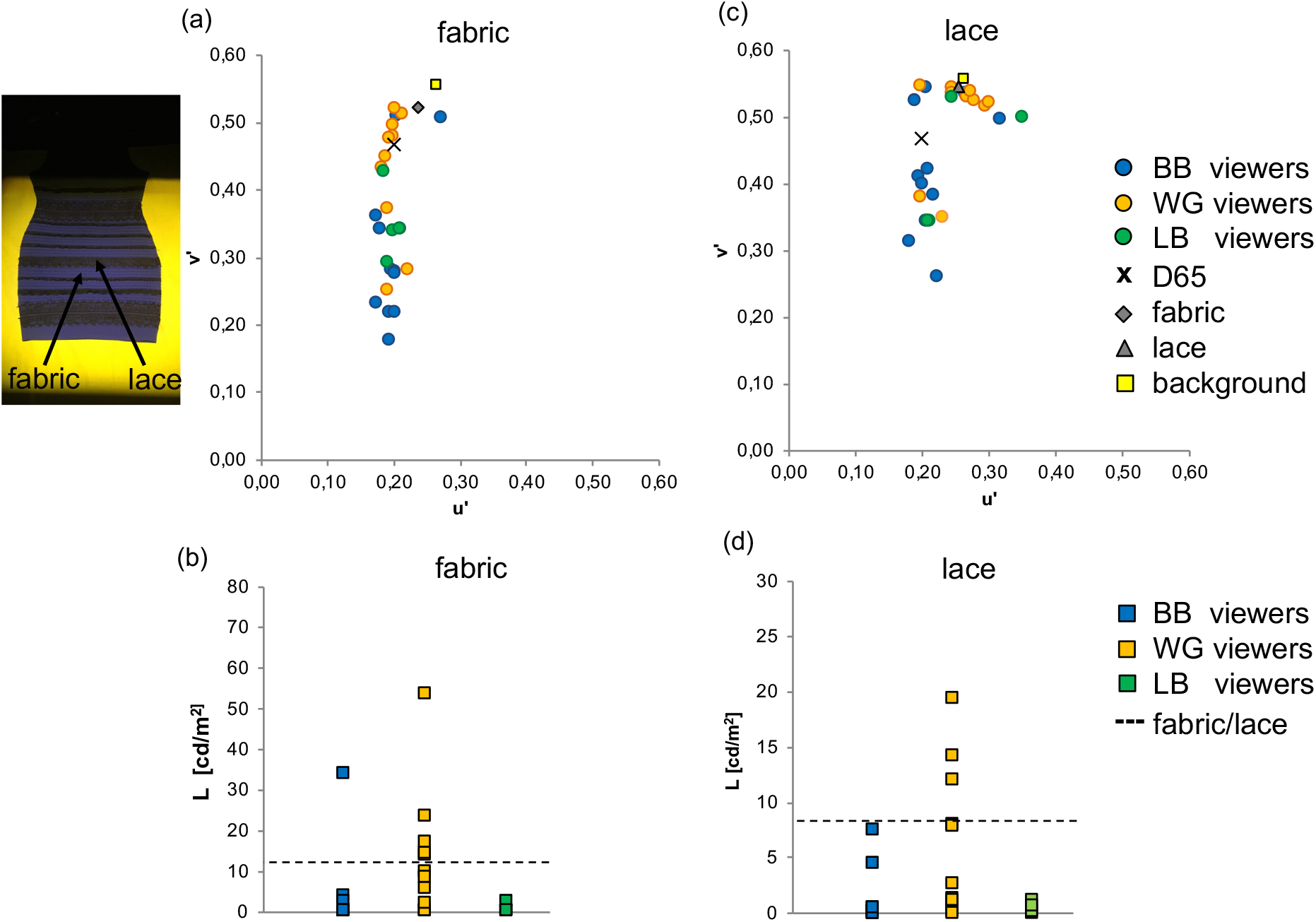
Settings of all observers for the appearance of the real dress presented on a yellow background cloth under projector light. (a) u’v’ chromaticities and (b) luminance of the matches for the fabric; (c) u’v’ chromaticities and (d) luminance of the matches for the lace, respectively. The colour locus of the background is indicated by the yellow square, the inserted image on the upper left shows the real dress on the yellow background, as presented in this experiment; arrows indicate the regions (fabric and lace) from which the measurements and colour matches were obtained. Yellow square marks the colour locus of the background cloth, otherwise same symbols as in figure 6.

The perceptual matches revealed that the chromatic settings for the fabric were again alined along the blue/yellow axis (figure 10). In addition, two clearly separated clusters were observed: one cluster was formed predominantly by WG viewers (WG: u’_mean_ = 0.195, v’_mean_ = 0.456; L_mean_ = 13.45) and was located in the achromatic/yellowish region between D65 and the colour locus of the fabric; the other cluster in the bluish region was formed predominantly by BB viewers (BB: u’_mean_ = 0.199, v’_mean_ = 0.308; L_mean_ = 3.78) and LB viewers (LB: u’_mean_ = 0.196, v’_mean_ = 0.350; L_mean_ = 0.67) viewers. Accordingly, the v’ and L values of the matches of WG viewers differed significantly from those of BB viewers (v’: p_WG-BB_ = 0.013; L: p_WG-BB_ = 0.016) and those of LB viewers (v’: p_WG-LB_ = 0.018; L: p_WG-LB_ = 0.026), respectively. Only two of the original 11 BB viewers were now located within the WG cluster, and one of the original WG viewers was now located within the BB cluster. Lightness matches of WG viewers were significantly higher than those of BB viewers (p_WG-BB_ = 0.016) and LB viewers (p_WG-LB_ = 0.026), respectively.

The ambiguity is also reflected in the perceptual settings for the lace: here, one cluster around the colourimetric locus of lace was formed predominantly by WG viewers (WG: u’_mean_ = 0.242, v’_mean_ = 0.508; L_mean_ = 6.18 cd/m^2^) and another cluster in the bluish region was formed mainly by BB viewers (u’_mean_ = 0.220, v’_mean_ = 0.420; L_mean_ = 1.20 cd/m^2^) and LB viewers (u’_mean_ = 0.254, v’_mean_ = 0.428; L_mean_ = 0.49 cd/m^2^). v’ coordinates differed significantly between WG and BB viewers (p_WG-BB_ = 0.023) and LB (p_WG-LB_ = 0.040), respectively, but not between BB and LB viewers (p_WG-LB_ > 0.1). Luminance settings of WG viewers were significantly higher than those of BB vierwers (p_WG-BB_ = 0.005), but not higher than those of LB viewers (p_WG-LB_ = 0.056).

##### Summary and conclusion of part 2

Overall, in both experiments using the real dress illuminated by blue&yellow projector lights, we found the chromatic settings for the dress (fabric and lace) distributed along the blue/yellow axis, and this was comparable to the pattern of distributions in the photograph. For the majority of WG and BB observers their original classifications as WG, LB and BB viewers were confirmed. Presenting the real dress on a black background did, however, not result in significant differences between the settings of WG, BB and LB viewers; in contrast, presenting the dress on the yellow background induced a clear perceptual ambiguity as known from the photograph. Likewise, WG viewers made higher lightness matches than the other groups, indicating an additional ambiguity in the lightness domain. Also, it is noticeable that the colour matches of almost all observers, including the WG viewers, showed (relative to the colorimetric locus of the dress fabric), a bias towards blue, which was more pronounced in the BB viewers than in the WG viewers, and more pronounced in the experiments involving the yellow background than the black background.

The reported perceptual shifts following the background change from “black” (or better to say “very dark yellow”) to bright yellow indicates a strong influence of the chromatic and luminance context on the colours of the dress. In the following, we investigated the influence of the background therefore more closely.

### 3. INVESTIGATIONS INTO THE ROLE OF THE BACKGROUND

#### 3.1. Effect of changing the background colour

Here we focused on the analysis of the matches made for the fabric and analysed the perceptual changes in chromaticity and luminance following the background change. The chromaticity matches made for the fabric with the black and the yellow background are plotted for direct comparison in figure 11.

**Figure 11.**
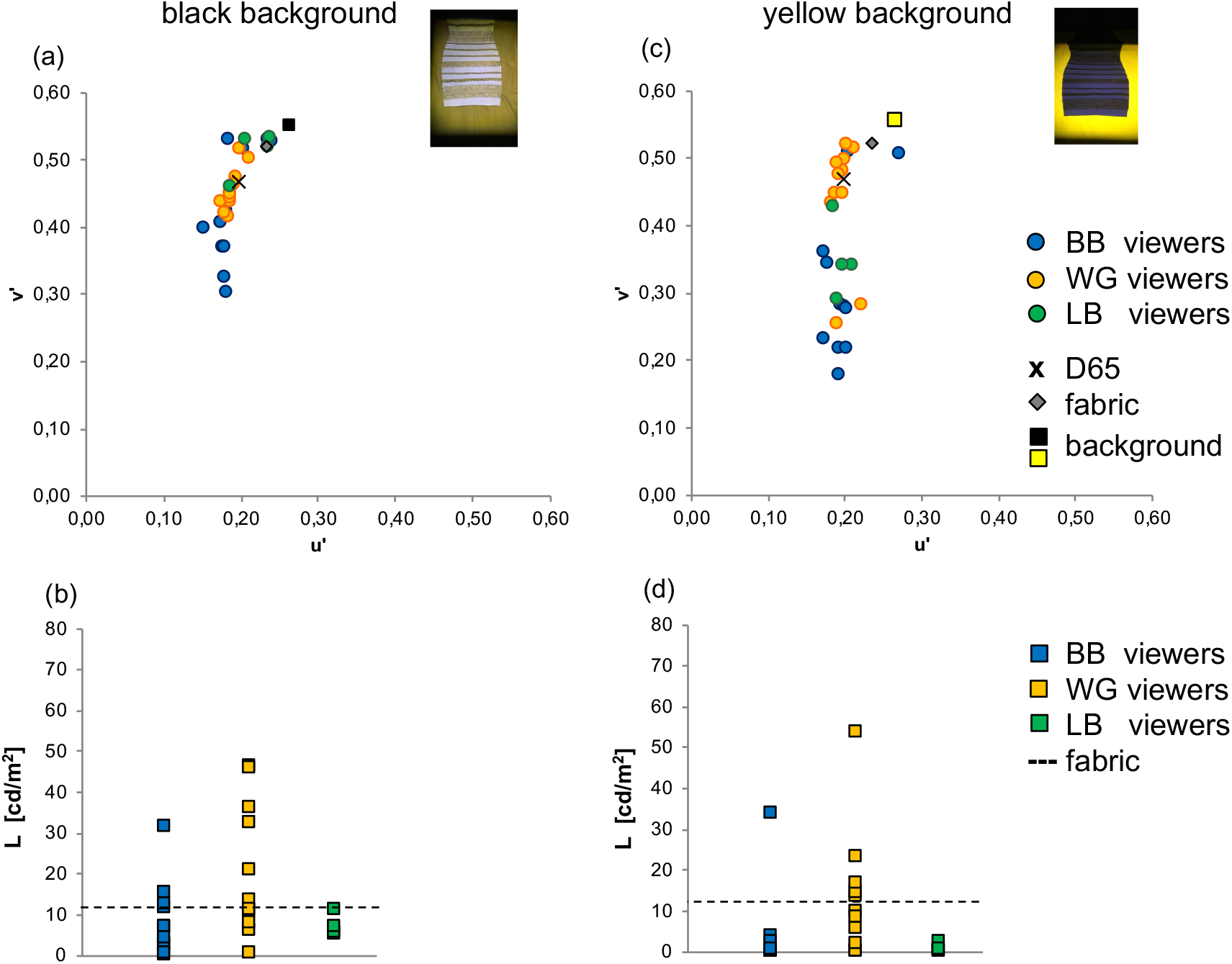
Effect of background change: colour matches made by the observers for the fabric of the dress on the black background (left) and on the yellow background (right). (a), (c) u’v’ chromaticities and (b, d) luminance of the matches. Same symbols as in figures 8 and 10.

Following the background change, the shift in chromaticity showed a significant difference between the WG viewers on one side and the BB and LB viewers on the other: it was found that the perception of WG viewers changed significantly less (WG: Δu’v’_mean_ = 0.04) than those of the BB viewers (BB: Δu’v’_mean_ = 0.12, p_WG-BB_ = 0.019) and LB viewers (LB: Δu’v’ mean = 0.16, p_WG-LB_ = 0.018); there was no difference between BB and LB viewers (p_BB-LB_ > 0.1). In other words, whereas the perception of the majority of BB and LB viewers shifted significantly towards a higher chroma of blue, the perceptual settings of the WG viewers remained clustered near D65.

Furthermore, all groups showed a reduction in the perceived luminance of the fabric (WG: ΔL_mean_ = 12.29 cd/m^2^; BB: ΔL_mean_ = 5.30 cd/m^2^; LB: ΔL_mean_ = 6.75 cd/m^2^), which was significant in BB viewers (p_BB_ = 0.007) and LB viewers (p_LB_ = 0.029), but not in WG viewers (p_WG_ > 0.1). Note that the black cloth and the yellow cloth differed mainly in their luminance, and only little in their chromaticities (see colour loci in figure 4). Not surprisingly, the higher luminance of the yellow cloth caused a lightness induction; what is surprising, though, is that the lightness change was significantly stronger in BB and LB viewers than in WG viewers and, furthermore, that there was an additional hue shift in the perception of BB viewers.

These findings indicate a significant difference regarding the influence of background on the different groups of observers. If this is correct, we should find BB viewers more affected by the absence of context information then the WG viewers, and the ambiguity should be reduced or lost. The following two experiments were designed to test this prediction.

#### 3.2. Effect of reducing the contextual background (masking experiments)

We tested the effect of chromatic context on the observers for the real dress and for the photograph. In the first experiment, we restricted the illumination of the real dress to the inner region of the skirt. Now, the viewers reported the colour of the dress fabric as white-beige or having a slight bluish tint, the lace was described as being gold (figure 12).

**Figure 12.**
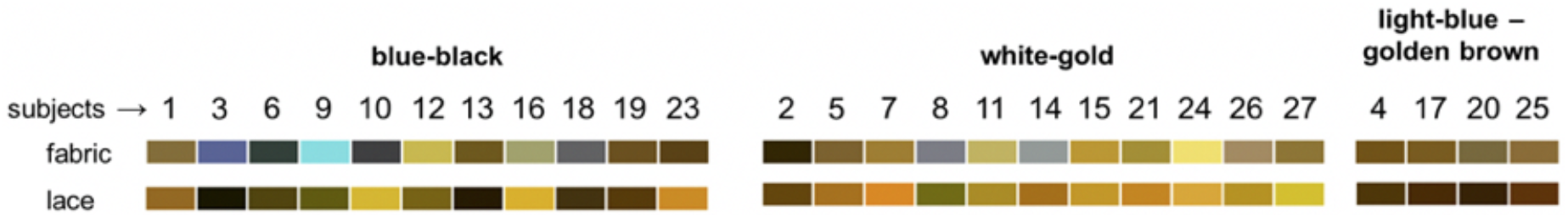
Appearance of the colour matches made by BB, WG and LB viewers for the real dress in the masked condition (i.e. illumination restricted to the inner region of the dress skirt).

The corresponding settings (figure 13) correlated closely with the uniformity of the colour names: the variation of the chromaticity settings u’ and v’ as well as in the lightness domain were strongly reduced; there were no significant differences between the perceptual groups (WG: u’_mean_ = 0.2168, v’_mean_ = 0.5070, L’_mean_ = 12.972 cd/m^2^; BB: u’_mean_ = 0.2030, v’_mean_ =0.4766; L_mean_ = 8.014 cd/m^2^; LB: u’_mean_ = 0.2370, v’_mean_ = 0.5281; L_mean_ = 5.308 cd/m^2^). Matches of all (but one) BB observers were now located around the fabrics locus in the yellow region and the achromatic region. In other words, the ambiguity was lost. Interestingly, the perceptual changes between seeing the entire dress and its masked version were significantly stronger in the group of BB viewers (Δu’v’_mean_ = 0.1698) and LB viewers (Δu’v’_mean_ = 0.1831), than in WG viewers (Δu’v’ = 0.0605; p_WG-BB_ = 0.008; p_WG-LB_ = 0.006). With respect to the luminance settings, the values of all groups were in the masked condition distributed around the actual (veridical) luminance of the dress, without group specific differences (WG: L_mean_ = 12.97 cd/m^2^, BB: L_mean_ = 8.01 cd/m^2^, LB: L_mean_ = 5.31 cd/m^2^); only BB viewers showed significantly higher luminance matches in the masked condition as compared to the full view condition (BB:ΔL_mean_ = 6.7 cd/m^2^; p_BB_ = 0.01).

**Figure 13.**
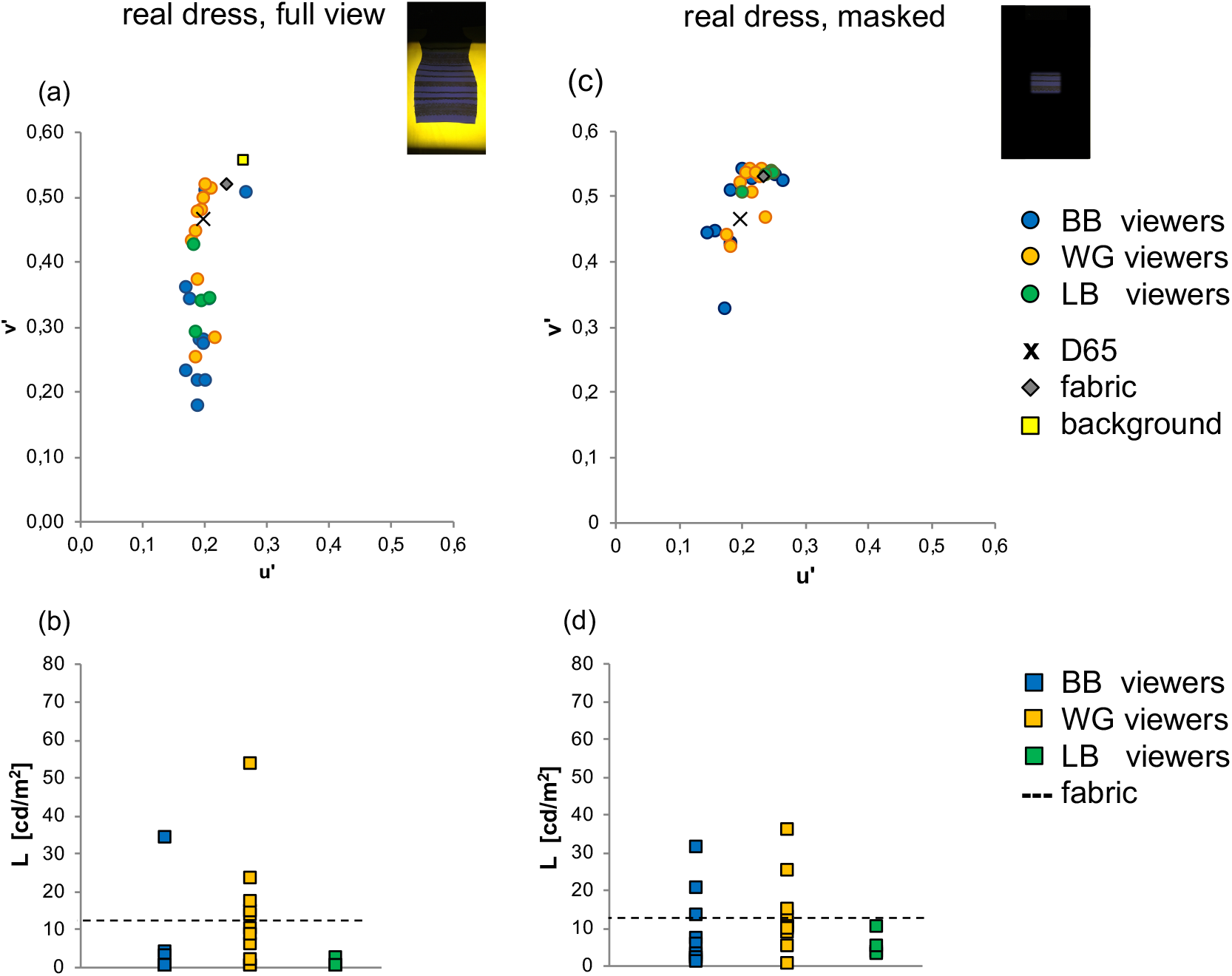
Effect of masking on the appearance of the fabric of the real dress. Left side: full view condition, right side: masked condition. (a), (c) u’v’ chromaticities and (b, d) luminance of the matches. Same symbols as in figure 10.

In the second experiment, the dress photograph was masked so that only two small patches (one for fabric, one for lace) were visible; again, in comparison to seeing the entire photograph, the colour appearance matches for the dress’ fabric became more uniform (figure 14); and as can be seen in figure 15, the variability of the settings was strongly reduced (WG: u’_mean_ = 0.1991, v’_mean_ = 0.4207; BB: u’_mean_ = 0.1959, v’_mean_ = 0.3894; u’_mean_ = 0.1981, v’_mean_ = 0.3754), but v’ coordinates still differed significantly between WG viewers and BB viewers (p_WG-BB_ = 0.016) and LB viewers (p_WG-LB_ = 0.026), respectively; there was no difference between BB and LB viewers (p_BB-LB_ > 0.1). In other words, the ambiguity was reduced but not completely lost. The change from full view to the masked condition also made BB viewers to increase their lightness matches significantly (ΔL_mean_ = 13.78 cd/m^2^, p = 0.001), while the lightness matches of WG viewers were significantly reduced (ΔL_mean_ = 31.73 cd/m^2^ p = 0.007), and LB viewers remained unchanged (ΔL_mean_ = 3.84 cd/m^2^, p > 0.1). Accordingly, the lightness matches of WG (L_mean_ = 31.73 cd/m^2^) and BB viewers (L_mean_ = 26.43 cd/m^2^) were significantly higher than the actual luminance of the fabric (WG: p_wg-fabric_ = 0.002, BB: p_BB-fabric_ = 0.004); only LB viewers (L_mean_ = 18.89 cd/m^2^) were close to the veridical luminance.

**Figure 14.**
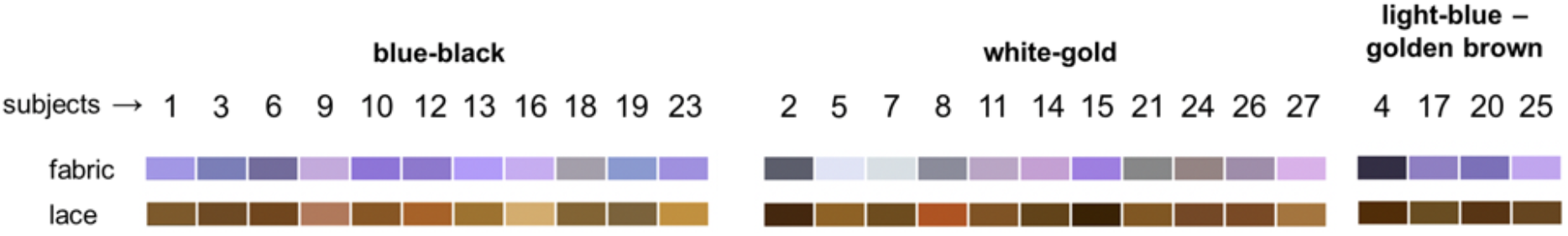
Appearance of the colour matches made by BB, WG and LB viewers for the fabric and lace of the photograph in the masked condition.

**Figure 15.**
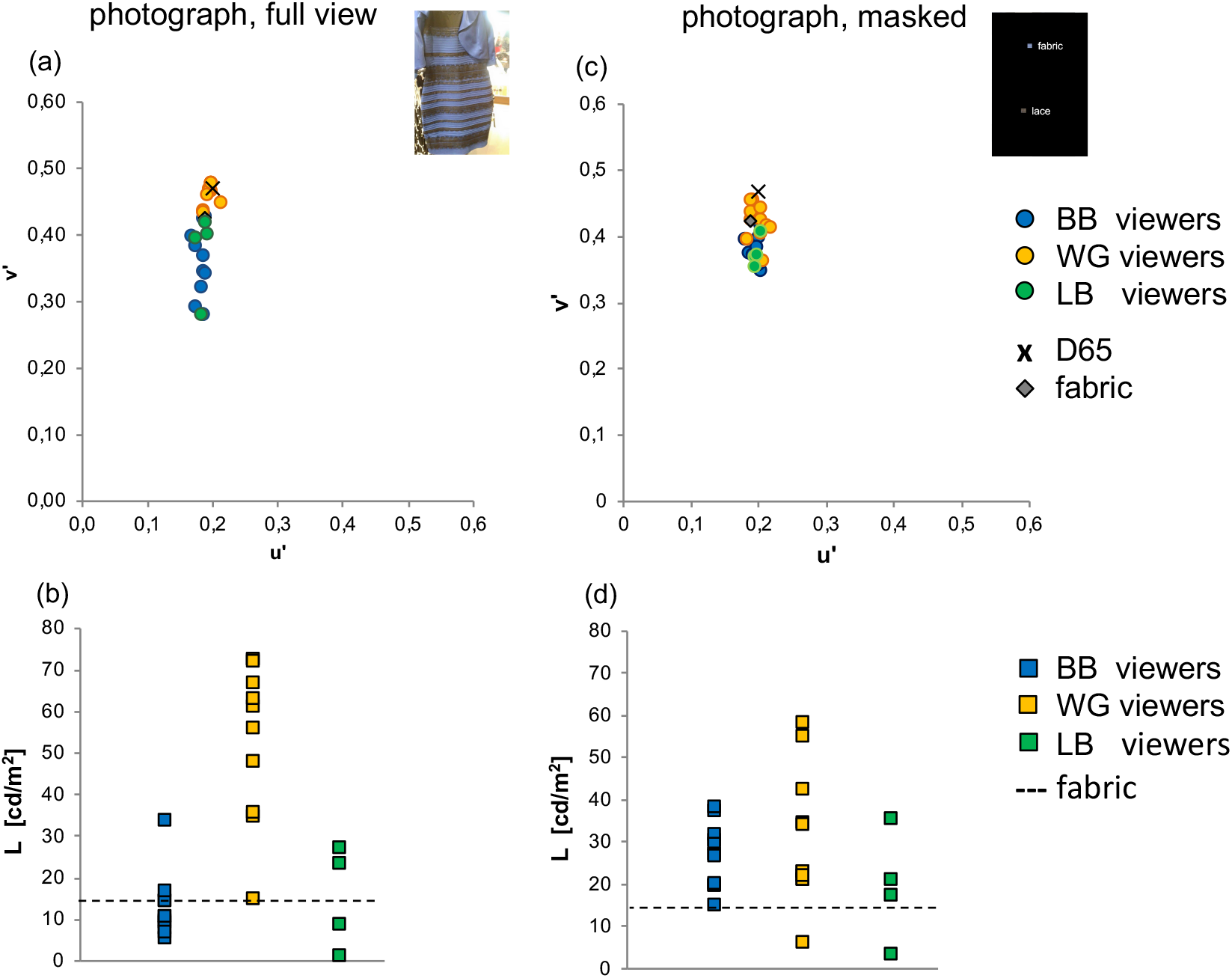
Effect of masking on the appearance of the dress’ fabric in the photograph. Left side: full view condition, right side: masked condition; (a, b) u’v’ chromaticties, (c,d) luminance of the matches; same symbols as in figure 6. Inserted images show the photograph in the full view (left) and in the masked condition (right); no text was present during the presentation of the masked photograph. Same symbols as in figure 6.

##### Summary and conclusion of part 3

The experiments of part 3 revealed (1) a strong influence of the chromatic context on the perception of the dress and (2) significant differences between the perceptual groups: for the dress fabric, the change of the background from black (i.e. dark yellow) to bright yellow induced in BB and LB viewers - but not in WG viewers - a reduction in lightness; also, BB and LB viewers perception shifted significantly towards blue, whereas WG viewers showed little to no shifts in chromaticity. It is interesting to note that the distinct blue shift of the BB viewers was induced by a higher luminance of the yellow background, not by a major change of its chromaticity. (3) Furthermore, in the mask experiments, the ambiguity was strongly reduced, whereby the settings of BB and LB viewers shifted significantly more than those of the WG viewers. Likewise, the lightness settings of BB and LB viewers changed significantly, while those of WG viewers did not. This is consistent with the findings from the background changing experiments and supports the notion that the perception of BB and LB viewers is more influenced by the background than that of the WG viewers.

### 4. THE DRESS UNDER CEILING LIGHT

Finally and as a control condition, we asked: how does the dress appear in a “normal” viewing condition? For that purpose, the real dress was now presented in a room fully lit by an artificial daylight-source on the ceiling. As before, the dress hung in front of the black background cloth and observers were asked to match the colours seen in the skirt part of the dress. The resulting appearance and the settings of all observers for the fabric as well as for the lace can be seen in figures 16 & 17.

**Figure 16.**
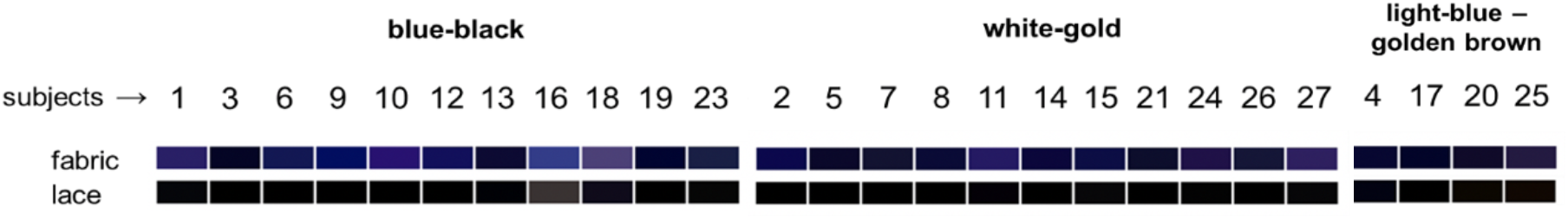
Appearance of the colour matches made by BB, WG and LB viewers for fabric and lace of the real dress under ceiling light.

**Figure 17.**
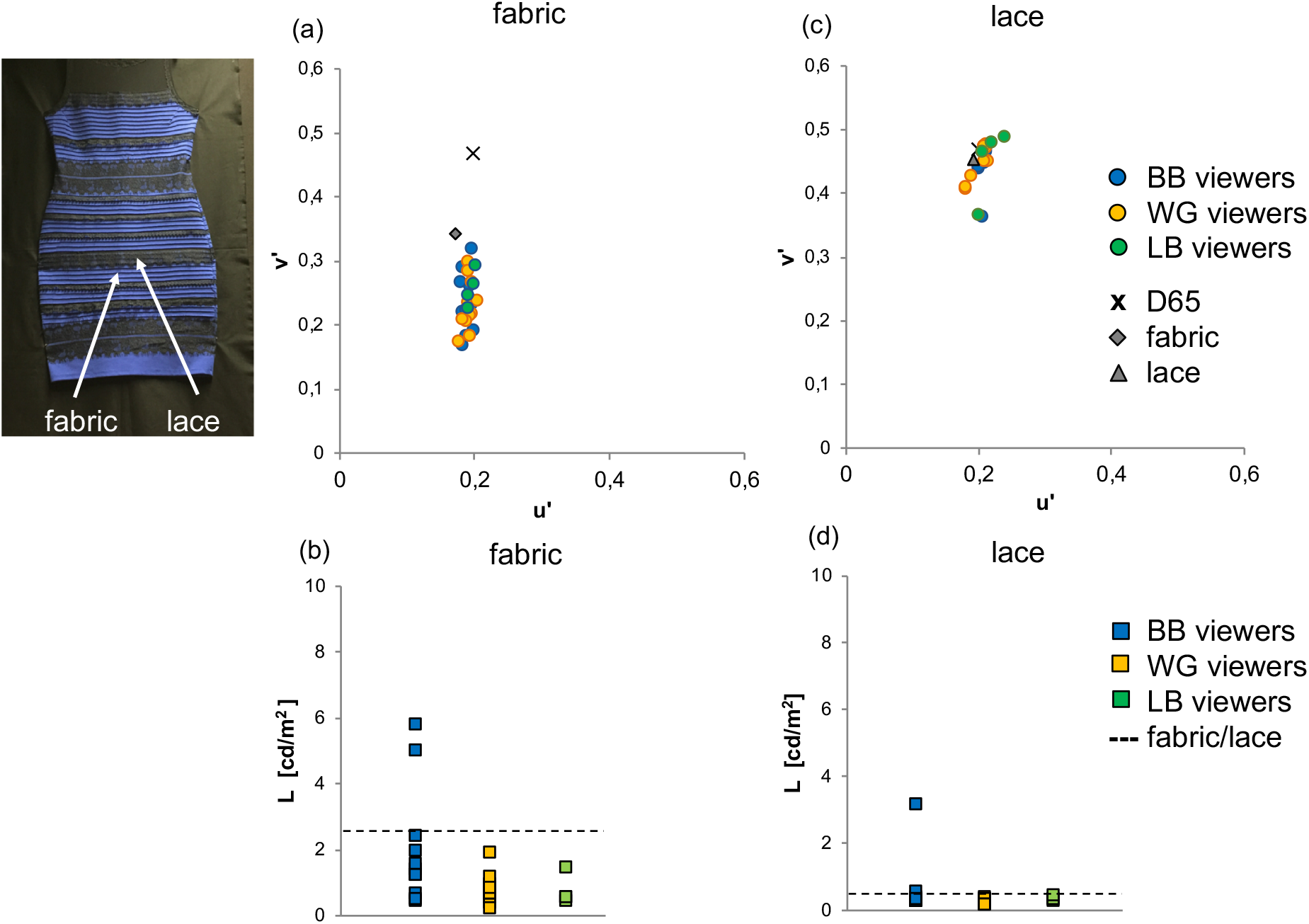
Observers’ settings for the appearance of the real dress presented on the black background in a fully lit room, under ceiling light. (a) u’v’ chromaticities and (b) luminance of the matches for the fabric; (c) u’v’ chromaticities and (d) luminance of the matches for the lace, respectively. The image on the upper left shows the real dress as presented in this experiment; arrows indicate the regions (fabric and lace) from which the measurements and colour matches were obtained. Same symbols as in figure 6.

There was still a variability noticeble along the v’ coordinate, corrresponding to a variability along the blue/yellow axis; there were however no group specific differences, i.e. WG, BB and LB viewers were evenly distributed within this cluster (WG: fabric: u’_mean_ = 0.1925, v’_mean_ = 0.2281; lace: u’_mean_ = 0.2037, v’_mean_ = 0.4516; BB: fabric: u’_mean_ = 0.1903, v’_mean_ = 0.2343; u’_mean_ = 0.2091, v’_mean_ = 0.4495; LB: fabric: u’_mean_ = 0.1965, v’_mean_ = 0.2560; lace: u’_mean_ = 0.2175, v’_mean_ = 0.4485. This corresponded to a choice of similar colour names to describe the colour of the fabric and the lace (namely „blue“, and „black“, respectively).

Similarly, the lightness matches for fabric and lace were close to the veridical values with no significant differences between the groups (figure 17 c,d): fabric: WG: L_mean_ = 0.80 cd/m^2^; BB: L_mean_ = 2.01 cd/m^2^; LB: L_mean_ = 0.73 cd/m^2^; lace: WG: L_mean_ = 0.19 cd/m^2^; BB: L_mean_ = 0.33 cd/m^2^; LB: L_mean_= 0.20 cd/m^2^). Thus, no ambiguity was observed when viewing the dress in a fully lit room, i.e., in a natural viewing condition; furthermore, when comparing qualitatively the matches for the fabric with the actual colour locus of the dress, a bias towards blue can be noticed in all settings; in other words, the observers saw the real dress as considerably more bluish than its colourimetric locus would predict.

#### Summary and conclusion of part 4

The settings for the dress illuminated by ceiling light within a fully lit room showed no ambiguity, although a slight variability of the settings was observed along the blue part of the blue/yellow axis. Furthermore, all observers showed a considerable blue bias in their perception of the fabric of the real dress, even in the “natural” viewing condition in a fully lit room. The lack of ambiguity here contrasts the perceptions reported by the same observers when viewing the dress under the projector light. Interestingly, knowing the “real colour” of the dress did not influence the classification of the observers as BB, WG and LB viewers when seeing the dress in the photograph again afterwards.

## DISCUSSION

In this study, we reproduced the scene of the dress photograph in our lab using the real dress and an artificial light situation and measured the corresponding colour perceptions of the observers. We found that the observers’ chromatic settings for the fabric of the real dress were distributed along the blue/yellow axis (figures 8, 10), with a variability that was comparable to viewing the dress photograph (figure 6); with the exception of only few observers, their previous classification as WG, BB and LB viewers held for their matches of the dress in the photograph and of the real dress. When the dress was presented on a yellow background (figure 10), the range of the distributions was substantially widened, due to a stronger blue shift (reaching higher saturation values) of the settings of BB viewers. Also, for the real dress, the range of settings of WG and LB viewers in the yellow colour region was enlarged owing to the more yellow colour locus of the fabric (again in comparison to the photograph). We conclude that our experimental setting with the real dress under our bi-colour projector light successfully reproduced a colour ambiguity similar to the one of the dress photograph. The ambiguity was specific for the particular experimental presentation and viewing situation of the dress, since no ambiguity was noted when the real dress was presented in a fully lit room under ceiling light (figure 17). In this latter condition, all observers described the colours of the dress as a vivid blue/black.

As a whole, the colour matches made for our dress photograph and the real dress are in accordance with the findings of previous studies of the dress photograph (e.g. [2, 13, 30]. Some of these studies, however, observed significant variations in lightness only [9, 13], whereas others report an additional ambiguity in the colour domain [2]; we found an ambiguity in both domains, colour and lightness, in the case of the photograph as well as the real dress. It has been argued that these differences may be attributable to the method of matching the dress colours; we used however the same method in both sets of experiments, and the cause for the differences remains therefore unclear.

All previous studies on the origin of the dress ambiguity have – to our best knowlege – used the dress photograph. Therefore, before we discuss our conclusion in the context of those studies, we need to emphasise an important difference between the different experimental approaches. For example, some studies on the dress photograph stressed the importance of top down influences, in order to cope with the sparse and ambigous information of that scene [2, 11, 19]: in particular, they showed how the colour of the dress in the photograph can be influenced by priming, i.e. by the previous exposure of the subjects to an unambigous and explicit appearance of the dress colour (white & gold or blue & black), produced either by induction [2, 19] or by colour constancy operations [11]. In contrast, our real scene did not contain ambigous information about the background or the lightfield in the scene as does the photograph, since observers were aware of the position of the light source and the nature of the background. This may impact the relative contribution of top down inferences and priors to colour constancy operations, as compared to the impact of image based processes, such as contextual colour computations. This is an important difference between the two experimental conditions and has to be kept in mind, when comparing, as in the following, our results with those of previous studies on the dress photograph. Furthermore, it means that we may have to look for additional, image based factors for explaining the ambiguity in our real scene. In the following we will discuss the importance of the visual context (background) for eliciting the ambiguity in our experiments and the findings from other studies on the dress photograph.

### Ambiguity depends on context

Our particular set-up enabled us to manipulate the surrounding context of the dress while quantifying the observers perceptual responses. Our results suggest a crucial role for the visual context (background) of the dress in eliciting the ambiguity, for the following reasons:

1. we found a strong influence of the background on the perceived colour and lightness of the dress, which was specific for the different perceptual groups: while the background change induced, in all groups, a reduction in lightness, it induced in BB and LB viewers, but not in WG viewers, an additional shift in chromaticity towards blue (figure 11).
2. in the masking experiments, the ambiguity was strongly reduced; importantly, the BB < viewers shifted their perceptual settings now towards those of the WG viewers, whose settings remained relatively unchanged (figures 13 & 15).

Taken together, these findings demonstrate that the settings of WG viewers were nearly independent of the presence or absence of the surrounding context, whereas BB ad LB viewers showed strong contextual influences. We conclude therefore that WG viewers preferentially compute the colour of the dress locally, within the dress region and with little reference to the background; the contextual colour processes of BB and LB viewers, on the other hand, seem to integrate signals across larger parts of the visual scene, including foreground (dress) and background.

This conclusion is supported by the findings of Toscani and colleagues [9] on the dress photograph who also reported differences in the processing of contextual cues between the perceptual groups. In their study, the contextual cues (additional heterochromatic patterns) were superimposed <u>onto</u> the dress and this influenced the WG viewers more than the BB viewers. Note that this does not contradict, but rather supports, our findings, since the contextual cues in our study refer to the background, whereas in the Toscani study, they refer to the local dress region. In fact, we expect WG viewers to respond more to the local luminance contrast than BB viewers. It would have been interesting to see how BB viewers respond to the same texture cues presented in the background, which unfortunately was not tested in the study of Toskani et al..

Finally, Dixon and Shapiro [31] proposed a colour constancy operation which discounts the effect of illumination by encoding visual signals at multiple spatial scales, and combining the information about illumination (low pass) and the object colour (high pass). When using this method on the dress photograph, they found that the size of the spatial filtering influences the colour of the dress: the smaller the size of the filters, the whiter became the dress. According to this hypothesis, we would expect WG viewers to use small scale filters, when looking at the dress and BB to use larger filters – which is consistent with the experimental findings in our study on the real dress.

Most similar to our masking conditions are the experiments by Jonauskaite and colleagues [32]. Using the photograph, they removed the dress background, leaving visible either a small patch or a vertical stripe of the dress, including fabric and lace. In the case of seeing exclusively the patch, the ambiguity was lost, but in their stripe condition, some of the ambiguity was still present, albeit strongly reduced, just as in our experiments of masking the photograph. We therefore agree on the conclusion of Jonauskaite et al. that - at least for the photogaph - some of the ambiguity can be attributed to the local dress texture or colours themselfs. At the same time, however, their findings, too, indicate a significant role for the background in the emergence of the ambiguity.

Manipulations of the chromatic context have also been used in studies using the dress photograph in order to prime observers to one or the other colour category: Drissi Daoudi and colleagues [19] changed the colour appearance of the dress by chromatic induction, i.e. by adding flancers next to it (white or blue); Witzel et al. [11] as well as Lafer - Souza et al. [33] have changed the appearant colour of the dress by manipulating the appearant illumination the photograph (dress in the shadow or in direct sun light). There is however a fundamental difference between ours and these studies: in the former studies the different surroundings did not directly produce an ambiguity, the ambiguity was only observed afterwards, after the observers were primed to the different colours of the dress (“one-shot learning” in the Drissi Daoudi study) or to the illumination (“illuminant prior” in the Witzel study) and then viewed the original photograph. In our study, in contrast, the very change from a black to a yellow background had already a specific and differential effect on WG and BB viewers, and increased the colour ambiguity. We conclude therefore that the contextual background is, at least in a real scene like ours, of crucial importance for the emergence of the dress ambiguity. In the following we will discuss the possible nature of the underlying contextual computations and their variability.

### The nature of the underlying contextual colour computations

When viewing an object in a real or realistic scene (e.g. a photograph), contextual colour computations involve local and spatially extensive processes, including multiplicative, v. Kries type adaptation and subtractive processes such as lateral inhibition [34-36]. The perceptual consequences of spatio-chromatic processes are twofold: 1) they can lead to local chromatic induction, a perceptual shift of the colour appearance away (colour contrast) or towards (colour assimilation) the inducing colour; and 2) they aid colour constancy, by encoding chromatic signals as ratios across large regions of the visual field [37, 38]; in addition, they can provide image information for an illuminant estimate (e.g. specular highlights, average chromaticity, shadows; [15, 39-41]). In the following we will discuss how these spatio-chromatic processes could account for the perceptual ambiguity observed in our experiments with the real dress.

In the background changing experiments, the shifts in the perceptual settings of the BB viewers for the dress’ fabric are consistent with an effect of chromatic induction, i.a. a shift towards blue as expected from a yellow inducer (the background cloth). Can we explain therefore our findings simply by individual variations in chromatic (and lightness) induction? Substantial individual variations have indeed been repeatedly reported for chromatic induction [19, 42-44] and can been explained by differences of the observers gaze or eye movements [43, 45, 46]. But neither ours nor other studies found evidence for a substantial difference in the viewing behaviour of the subjects when viewing the dress (figure S1, suppl.), or other scenes [19, 43]. Furthermore, testing the individual variability in the amount of chromatic induction of WG and BB viewers in another set of independent experiments using a centre surround paradigme, we found no evidence for stronger induction effects in BB viewers than in WG viewers (figure S2, suppl.).

With respect to colour constancy operations, the spatio-chromatic processes require integration across large parts of the visual field, in order to avoid effects of local induction. Different results can be expected if the contextual processes include different regions of the scene. Our findings therefore indicate group specific differences in the contribution of the background: the contextual effects seen in BB and LB viewers are consistent with long-range contextual colour operations, such as ratio taking computational steps (low to mid level processes) or an image based estimation of the illumination; the data of the WG viewers, on the other hand, indicate a higher weight of the colour computations on the dress region (foreground) than on the background, and therefore are determined predominantly by the chromaticities of the dress region.

We have to ask therefore: what are the factors determining the operating range and the selection of regions for these processes, and what is the origin of their individual variations?

### Perceptual organisation and segmentation hypothesis

The range of contextual processes can be strongly influenced by the perceptual organisation of a scene. This is demonstrated by the effect of perceptual organisation on colour and in particular lightness phenomena, like for example the Koffka ring [47] or Whites illusion [48, 49], where the segmentation or grouping of surfaces determines their lightness/colour. The perceptual organisation itself is controlled by objective as well as subjective cues: objective, cues are based on image content such as luminance contrast, edge classification, form and texture; subjective cues come from cognition such as Gestalt rules, i.e. the belongingness to a figure or background and, importantly, the interpretation of the light field [16, 37, 48, 50, 51].

The visual light field in particular deserves attention in the context of lightness and colour constancy: by visual light field we mean the perceived direction, chromaticity and brightness of the light illuminating objects in a visual scene [41]. Due to the effects of e.g. shadowing, transparency and interreflexions, the light field in three dimensional scenes is inherently complex and inhomogeneous [52, 53]. Lightness and colour constancy operations need to take this into account and require a segmentation of the scene into regions of approximately uniform illumination, otherwise the contextual processes would be detrimental for achieving a robust percept. It therefore makes sense that the perceptual organisation should control the spatial extend of colour operations, by grouping surfaces which share a common illumination, and segmenting regions with different illuminations; such regions have been termed frames, windows or layers [54-58]. Studies on colour constancy showed that the underlying contextual processes are indeed controlled by the spatial luminance structure of a visual scene (texture, spatial frequency, and depth plane) and in effect organise the visual scene into “illumination frames” (computational units for chromatic adaptation and colour constancy [35, 36, 59, 60]. In particular, it was demonstrated that segmentation by depth plane supports colour constancy in scenes with heterogenous illumination [60].

The results of our background changing experiment (experiment 2) are an indication for the relevance of the luminance structure in our visual scene for the extend of spatio-chromatic processes: here, the observer specific chromatic shifts were produced by a change in the luminance structure of the scene (change of the background from black to bright yellow), with little accompanying chromatic change. It has been shown for chromatic induction that high luminance contrast inhibits spatio-chromatic interactions [61] and it can therefore be expected that particular levels of luminance contrast in combination with the individual contrast sensitivity of an observer influences the segmentation of the scene. Indeed, Dixon and Shapiro [31] reported that WG viewers have a higher sensitivity for luminance contrast (in particular at low spatial frequencies) than BB viewers. This could explain why WG viewers are more prone to a segmentation of dress and background than BB viewers.

We propose therefore the following “segmentation hypothesis” as an explanation for the ambiguity of the dress. This hypothesis assumes a different perceptual organisation of the visual scene by WG, and BB viewers, respectively, as it is despicted in figure 18: based on the individual response to the luminance structure, the scene of WG viewers is segmented into several, separate frames (dress frame and background frame) or into one, “global” frame (dress+background frame), as the case in BB viewers. Accordingly, the contexual computations (induction and/or illumination estimate) are either restricted to the dress region (WG) or extend into the background (BB viewers) and this will consequently result in different colour percepts. In addition, in WG viewers, strong contextual interactions between lace and fabric within the dress region can be expected to further enhance their white/gold percept.

**Figure 18.**
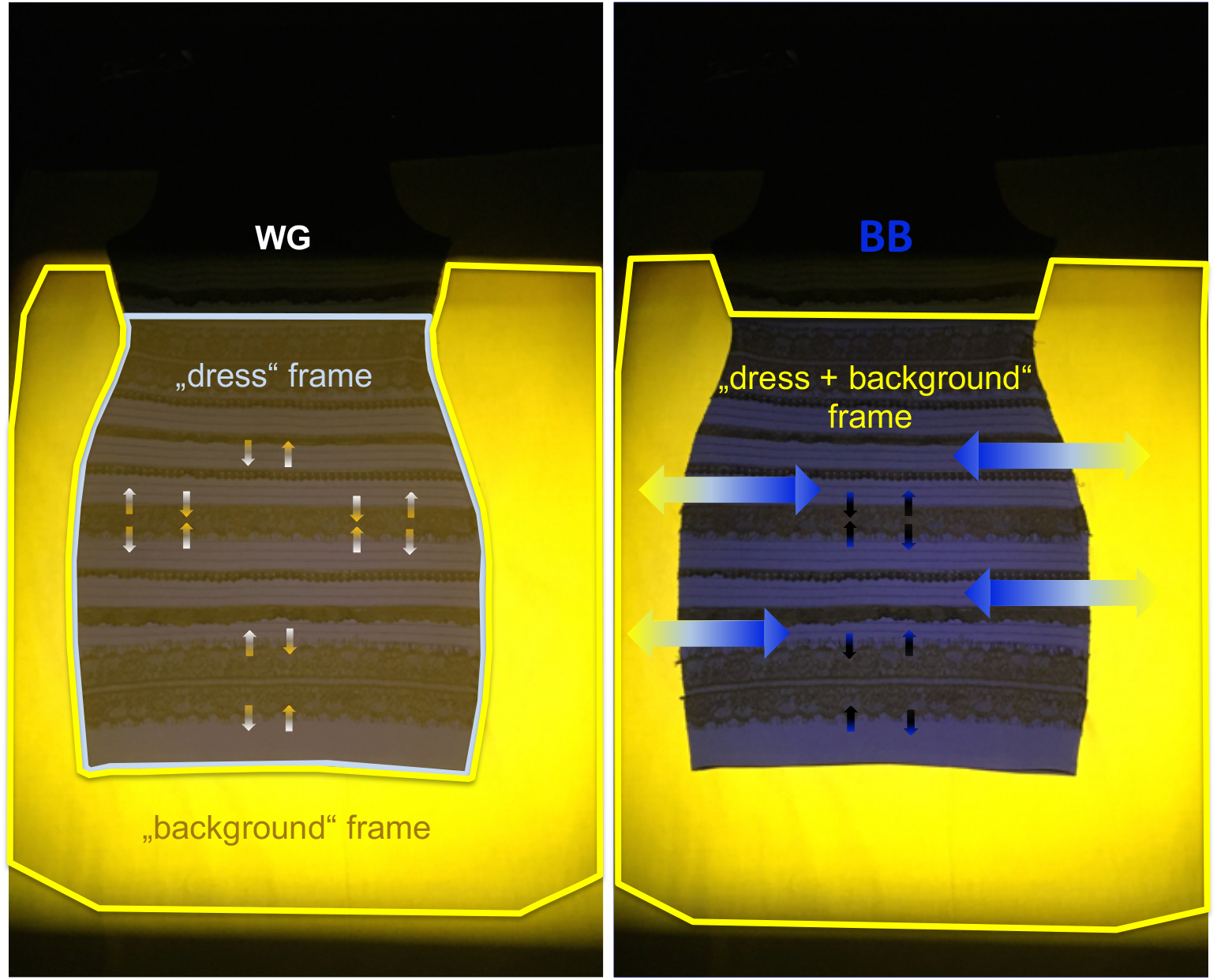
Segmentation hypothesis: a simplified sketch of the proposed perceptual organisation of the dress’ visual scene and the contextual interactions. Light blue and yellow lines delineate spatial regions for colour computations of WG viewers (left; two regions, a “dress” frame and a “background” frame) and of BB viewers (right; one region, the combined “dress + background” frame); arrows symbolise contextual interactions between fabric and lace (small arrows) and between the dress and its background (large arrows).

Similarly, the segmentation hypothesis can also be applied to the dress photograph. Here, the reported ambiguity of the visual light field may not only result in different illuminant estimates [9, 62] but also in a different segmentation of the scene during colour constancy operations: most of the BB assume the illumination in the scene of the photograph to come from the front and/or above the dress, in other words, they assume a uniform light field; WG viewers on the other hand, interpret the bright background as light coming from behind the dress, and the dress in the shadow, i.e. they assume two light zones with different properties. In short, the observer specific interpretations of the visual light field predict differences in the segmentation of the scene and consequently different operating ranges of the colour computations. With respect to colour constancy, this would mean that WG viewers compensate the local illumination on the dress region, and the BB viewers the global illumination across the entire scene.

### Who sees the colour of the dress veridical?

One last question remains: who saw the colours of the dress „correctly“? Our answer: neither of our group of observers: in the experiments with the bicolour projector light, the settings of WG observers were close to the actual colourimetric locus of the real dress; on the other hand, BB viewers saw the dress correctly with respect to its appearance in the “natural viewing” conditions, i.e. in scenes with access to contextual information. In other words, WG viewers were more veridical than BB viewers with respect to the local sensory input, but BB viewers were veridical with respect to the object colour of the dress.

## SUMMARY AND CONCLUSION

This study shows that colour ambiguity, previously thought to be specific for an artificial scene, can be reproduced in a real scene. We find group specific differences in the relative contribution of the local dress region (foreground) and the surrounding background to the computation of the dress colour (WG, BB, LB), which account for the colour ambiguity.

We interpret our findings within a framework of perceptual organisation for colour constancy: namely, that WG observers compute the dress colour based on local information from the dress region, whereas BB viewers process the dress colour with information from the spatial context of the scene. Our segmentation hypothesis proposes that the luminance structure (in our real scenes) and the interpretation of the light field (in the case of the photograph), segment the visual scene, and affect the range of contextual colour computations, including colour constancy operations. Individual differences in the observers segmentation processes ultimately result in the observed color ambiguity. Additional inferences from memory or assumptions about properties (color and lightness) of the illumination can also potentially influence the perceived colours either directly or indirectly via the perceptual organisation and the computation of illuminant estimates.

Thus, our segmentation hypothesis does not contradict existing explanations of the colour ambiguity regarding cognitive inferences, but it extends them by adding perceptual organisation and scene segmentation as an additional source for individual variability and the possible origin of colour ambiguity. Future studies on phenomena like the dress illusion are important in order to understand the role of perceptual organisation in colour constancy and the relative contribution of top-down inferences and image based cues.

## Acknowledgements

We thank your test subjects for their patience and diligence in matching the colours of the dress. The research was supported by the Excellence Initiative of Tuebingen University and the Tistou & Charlotte Kerstan foundation.

