## Supplementary figures S1 & S2 for "Colour Ambiguity in Real Scenes and the Role of Perceptual Organisation"

### SUPPLEMENT

#### 1. Measurement of observers' gaze when viewing the dress

In a separate set of experiments, viewing behaviour of another group of observers was recorded when inspecting visually the scene of the real dress on the yellow background in an independent set of experiments. Observers were catagorized according to their verbal statements (self description as BB, WG or LB viewer) when seeing the original dress photograph. Eye tracking (SMI Eye tracker) was used to find out whether BB observers (n = 11), WG observers (n = 11) and LB observers (n=8) differed in their viewing strategies and time spent looking at the background or the dress. The dress was presented hanging against the same yellow background as used in the experiments described above, on the backwall of an otherwise black light box (1x1x1m); the scene was illuminated from above by adjustable LEDs emitting "sunlight". The subjects were instructed to "look around in the scene" for one minute, while it was tracked where and how long they directed their gaze to the dress or the yellow background. For this purpose, the dress and the background were defined as separate AOI's (Areas of Interest) and the total time in which the test person directed the eyes to the dress or the background was evaluated. Region specific gaze time was analysed using semantig mapping ("BeGaze" software (SMI)). Statistical analysis was performed using a single factor ANOVA with normal distribution of the data. A Shapiro-Wilk test was used to check the normal distribution. The average time spent by BB, WG and LB viewers on the dress or the background is displayed in figure S1: all groups spent more time looking at the dress than at the background (BB: dress: 46908 ± 4494 SEM [ms], background: 9794 ± 4225 SEM [ms]; WG: dress: 48914 ± 4621 SEM [ms], background: 8005 ± 4057 SEM [ms]; LB: dress: 51978 ± 4232 SEM [ms], background: 5562 ± 3659 SEM [ms]; there were, however, no group specific differences between the gaze times (dress: WG vs. BS: $p = 0.69$; WG vs. BB: $p = 0.50$; BS vs. BB: $p = 0.097$) (background: WG vs. BB: $p = 0.59$; BS vs. BB: $p = 0.13$).

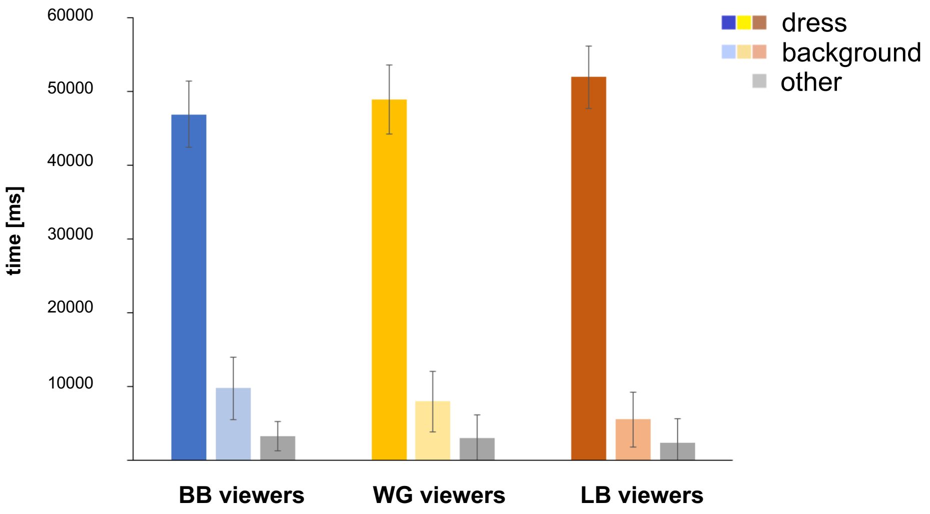


**Figure S1.** Time spent by observers of the different perceptual groups on average for viewing the dress or the yellow background; different colours refer to the different perceptual groups. The category "other" denotes time left to blinking or closing briefly the eyes.

### 2. Induction by the yellow background

Chromatic induction was measured in an independent experiment with another group of subjects. Twelve BB and twelve WG observers took part in the experiments, they were catagorized according to their self description when seeing the original dress photograph. Achromatic settings were obtained using a centre-surround paradigme, whereby a small central disk (diameter 0.9°; u' = 0.1978 v' = 0.4658 , L = 17 cd/m$^2$) was surrounded by an inducing ring (diameter 9.2°, u' = 0.2623, v'= 0.5570, L = 22 cd/m$^2$); the inducer had the same chromaticities as the yellow background cloth in the experiments with the real dress. Induction was measured by a nulling procedure, whereby subjects had to adjust the achromatic appearance of the central test field. The effect of induction by a yellow surround can - in general – be expected to shift the appearance of the central testfield towards blue, corresponding to shifting the subjective achromatic colour locus away from D65 towards yellow. The latter can indeed be seen in figure S2, but importantly, there was no systematic difference between the achromatic settings of BB and WG viewers (Tukey's test, BB vs WG: p(u') = 0.50, p(v') = 0.236, p(L) = 0.90).

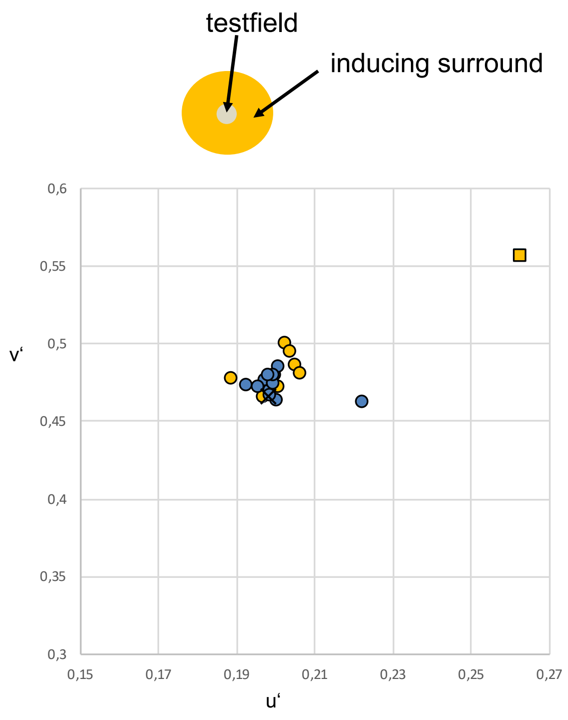


**Figure S2.** CIE 1976 UCS chromaticity diagram: achromatic settings of WG (yellow dots) and BB viewers (blue dots) for a small disk surrounded by a yellow ring (yellow square); x denotes the colour locus of D65. The inserted figure shows the stimulus arrangement.
